# Molecular fingerprints in the hippocampus of alcohol seeking during withdrawal

**DOI:** 10.1101/2023.08.24.554622

**Authors:** Roberto Pagano, Ahmad Salamian, Edyta Skonieczna, Bartosz Wojtas, Bartek Gielniewski, Zofia Harda, Anna Cały, Robbert Havekes, Ted Abel, Kasia Radwanska

## Abstract

Alcohol use disorder (AUD) is characterized by excessive alcohol seeking and use. Here, we investigated the molecular correlates of impaired extinction of alcohol seeking using a multidimentional mouse model of AUD. We distinguished AUD-prone and AUD-resistant mice, based on the presence of ≥ 2 or < 2 criteria of AUD and utilized RNA sequencing to identify genes that were differentially expressed in the hippocampus and amygdala of mice meeting ≥ 2 or < 2 criteria, as these brain regions are implicated in alcohol motivation, seeking, consumption and the cognitive inflexibility characteristic of AUD. Our findings revealed dysregulation of the genes associated with the actin cytoskeleton, including actin binding molecule cofilin, and impaired synaptic transmission in the hippocampi of mice meeting ≥ 2 criteria. Overexpression of cofilin in the polymorphic layer of the dentate gyrus (PoDG) inhibited ML-DG synapses, increased motivation to seek alcohol and impaired extinction of alcohol seeking, resembling the phenotype observed in mice meeting ≥ 2 criteria. Overall, our study uncovers a novel mechanism linking increased hippocampal cofilin expression with the AUD phenotype.

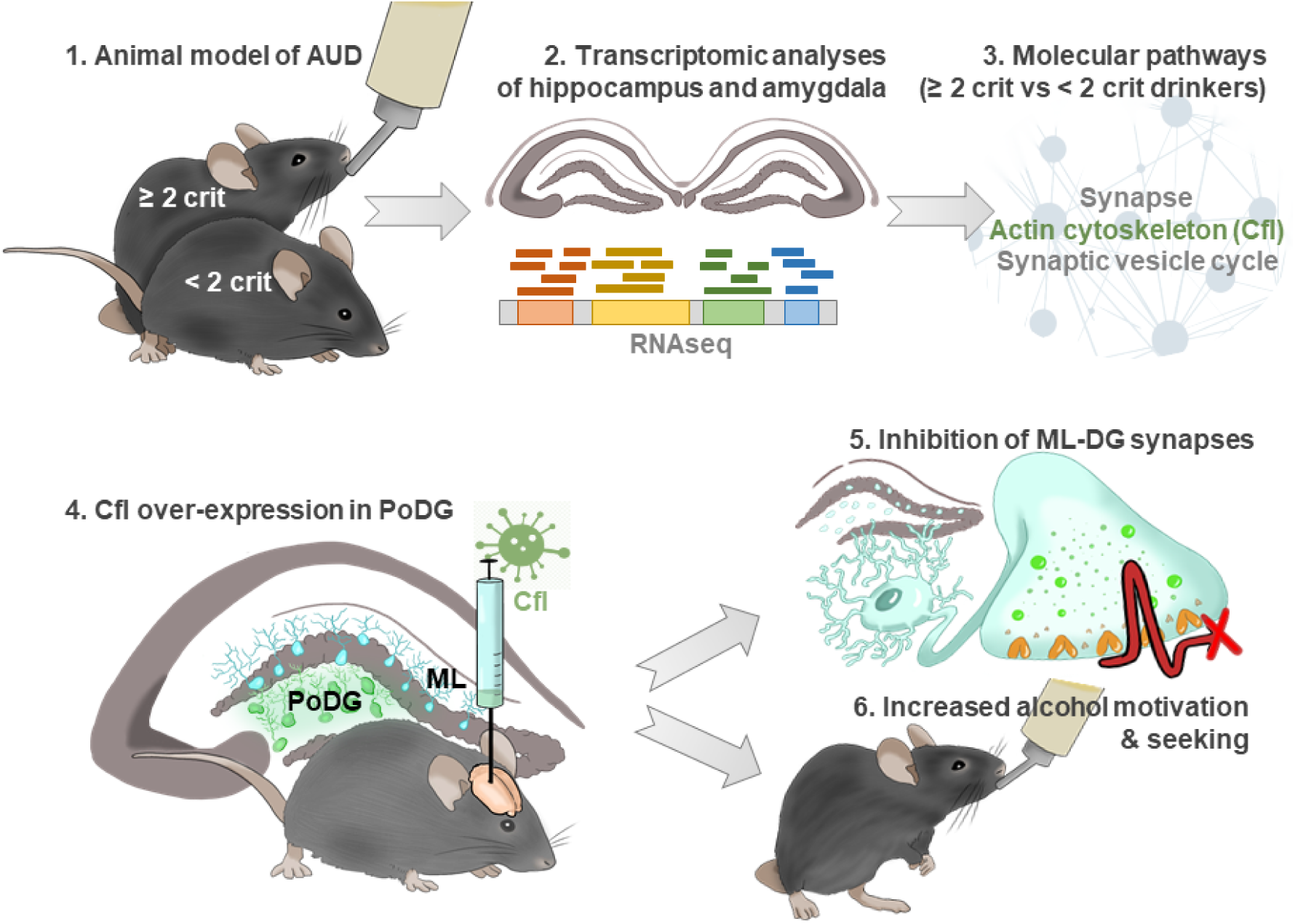

## INTRODUCTION

Alcohol use disorder (AUD) is a progressive and debilitating psychiatric disease characterized by pathological alcohol craving and motivation to consume alcohol, as well as cognitive rigidity. This condition results in an excessive focus on alcohol procurement and alcohol use in daily routines ^1^. Although AUD is one of the leading causes of premature deaths globally, pharmacological interventions aiming to control alcohol misuse are limited, with significant negative side effects, and therfore infrequently prescribed and used ^2–4^. To identify a new therapeutic approach to battle AUD, a neurobiology of the disease must be elucidated. So far most of the molecular studies focused on the quantitative aspects of alcohol misuse ^5^-several candidate genes and molecular pathways that affect the amounts of consumed alcohol both in humans and animals have been identified ^6–9^. Moreover, in recent years an accumulating number of studies focus on the biology of complex alcohol-related behaviors, such as compulsivity ^10–14^, cognitive inflexibility ^15^, or choice between alcohol and natural rewards ^16–18^. Still, the molecular processes that affect behavioral hallmarks of AUD beyond alcohol consumption remain poorly understood. To develop a successful prevention and therapeutic control of AUD progression, the neuronal basis of all AUD-related behaviors must be recognized.

Here we focused on the molecular correlates of excessive alcohol seeking induced in the alcohol-predicting context during withdrawal. Such behaviour reflects individual focus on alcohol procurment and use, as well as cognitive inflexibility characteristic for AUD patients ^1, 19^. Toward this end we developed a mouse model of AUD that has been pharmacologically-validated ^20–22^ and proven to have significant translational value ^23–25^. The model is based on four DSM-5 criteria of the disease ^1, 20^. (i) Craving, or a strong desire or urge to use alcohol. We measured motivation to obtain alcohol in a progressive-ratio schedule. (ii) The subjects spents a great deal of time in activities necessary to obtain alcohol. We measured the extinction of alcohol seeking during periods of forced abstinence. (iii) The subject takes alcohol in larger amounts than intended. We measured alcohol intake during alcohol relapse after abstinence. (iv) Unsuccessful efforts to control alcohol use. We measured alcohol seeking induced by alcohol-predicting cues and during signalled periods of alcohol non-avaliability. The model allowed us to distinguish the animals that exhibit consistent AUD-prone phenotype, as they were positive (uppermost 35% of the population) in at least two AUD-related tests (≥ 2 crit mice), and the AUD-resistant mice that were positive in none or one test (< 2 crit animals). Next, we used the new generation RNA sequencing (RNA-seq) to characterize differentially expressed genes in the hippocampus and amygdala of ≥ 2 crit and < 2 crit animals seeking for alcohol during alcohol withdrawal. We focused on the hippocampus as aberrant hippocampal synaptic plasticity is causally linked with cognitive and motivational aberrations characteristic for AUD, including drug seeking induced by drug-predicting contexts and cues ^21, 22, 26–30^. In particular, the manipulations that ablate adult neurogenesis in DG increase drug consumption and motivation to seek for drugs, as well as invigorate drug seeking induced by associated cues and contexts ^29, 30^. Moreover, the CA1 area and subiculum have been implicated in drug-induced place preference and context-induced alcohol and drug seeking ^26–28, 31–34^. On the other hand, the amygdala has been identified as a key region of the neural circuits implicated in the regulation of incentive salience of alcohol- and drug-associated cues, as well as cue-induced reinstatement of drug seeking ^23, 24, 35–41^. Furthermore, the amygdala was implicated in regulation of alcohol consumption despite negative consequences and alcohol choice over natural rewards ^11, 16^. The molecular processes that underlie the functions of the hippocampus and amygdala in AUD are still largely unknown.

Our data showed that the variance in the transcriptome between the < 2 crit and ≥ 2 crit drinkers after alcohol withdrawal primarily involves hippocampal genes related to the cytoskeleton and synaptic function, including actin binding molecule, cofilin (Cfl) ^42^. Accordingly, *ex vivo* electrophysiology was used to characterize pre- and post-synaptic changes in the hippocampus of the < 2 crit and ≥ 2 crit mice. Finally, we investigated the role of the hippocampal Cfl in the AUD pathology using the local expression of Cfl delivered by adeno-associated viral vectors. Overall, our study identifies transcriptomic differences between the AUD-prone vs -resistant drinkers during alcohol withdrawal. We also describe a novel mechanism that links Cfl-regulated synaptic plasticity in the hippocampus with AUD phenotype characterized by high motivation to seek for alcohol and impaired extinction of alcohol seeking during withdrawal.

## RESULTS

### Characteristics of AUD-prone and -resistant mice

To identify AUD-prone and -resistant mice we used a mouse model of the disease in the social context of IntelliCages ^20^. C57BL/6J mice (n = 58) went through a long-term training consisting of the introduction of alcohol (4, 8, 12%, days 1-12) and alcohol free access period (FA, 10%, days 13-47). During the 4-12% and FA mice had unlimited access to alcohol in the reward corner. Alcohol availability was signaled by the cue light presented each time a mouse entered the corner and each nosepoke in the corner gave access to alcohol for 5 seconds (fixed ratio 1, FR1). Next, we assessed behaviors that resemble DSM-5 criteria for AUD ^1, 20^: high motivation to drink alcohol was measured as a number of nose-pokes in the reward corner performed in a progressive-ratio schedule of reinforcement test when mice had to make an increasing number of nosepokes (FR2, 4, 8, 12, 16, 20, 24, 28…) in order to get access to alcohol for 5 seconds (Motivation); excessive alcohol seeking was measured as number of nosepokes in the alcohol corner when the corner was inactive and nosepokes had no programmed consequences (Extinction); reactivity to alcohol-predicting cues was assessed as nosepokes in the alcohol corner during presentation of the cue light when alcohol was not available (Cue relapse) ^43^; lack of control over alcohol consumption was assessed as alcohol consumption (g/kg/day) when the alcohol corner was activated after withdrawal (Alcohol relapse); while lack of control over alcohol seeking was measured as the change of nosepokes number to the alcohol corner during the non-active vs. active phases of the test (Persistence) (**Figure** 1A). AUD score was calculated as a sum of normalized scores from all AUD tests, and AUD index as a sum of positive results (top 35%) in all tests ^20, 44^. Mice were distinguished based on the DSM-5 criteria^1, 20^: AUD-prone drinkers were positive in at least two AUD tests (AUD Index ≥ 2 crit), AUD-resistant drinkers were positive for none or one criterion (AUD Index < 2 crit). Overall, 38% of the mice were indicated as AUD-prone drinkers (**Figure** 1B).

**Figure 1.**
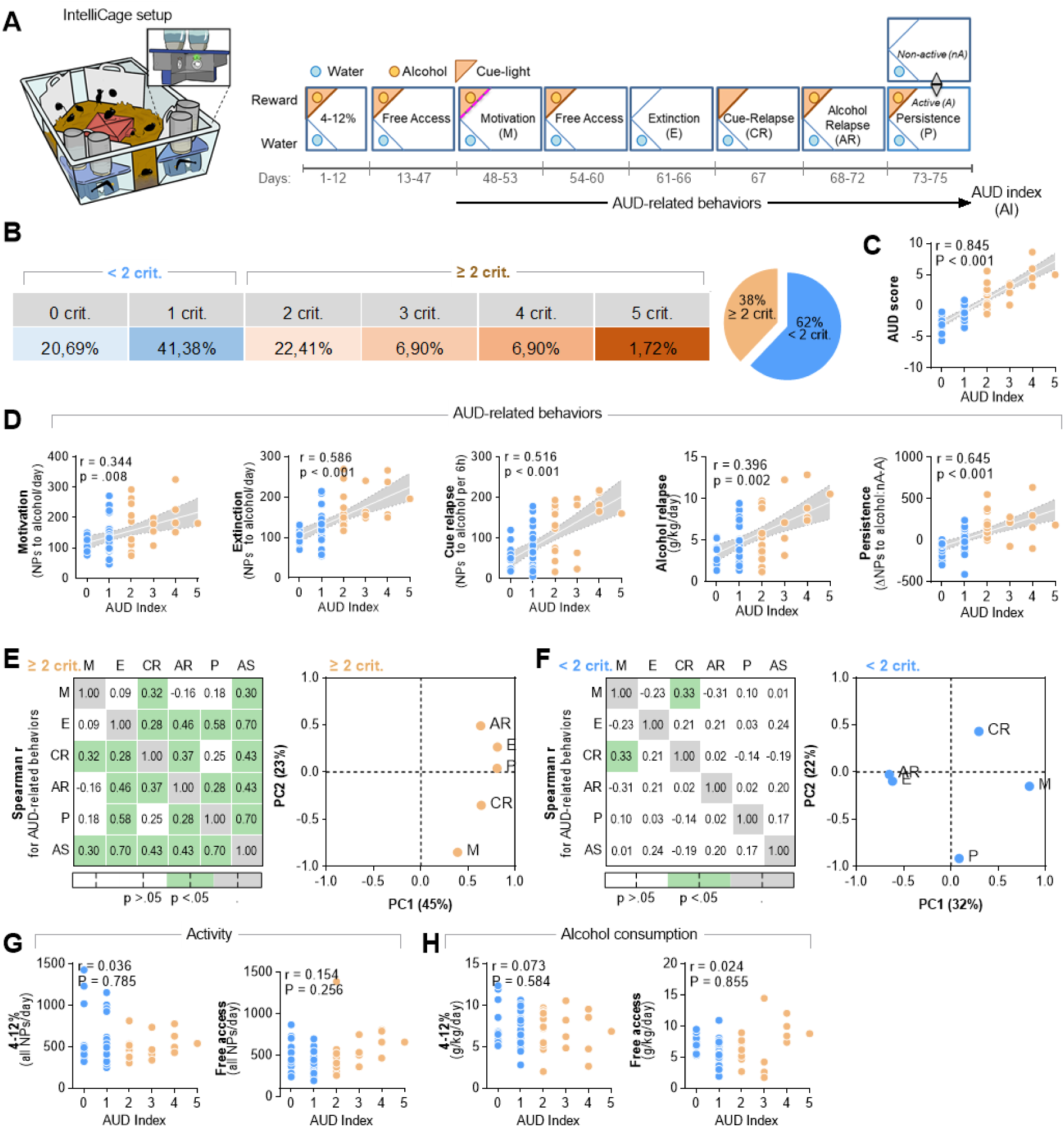
AUD-related behaviors of the < 2 crit and ≥ 2 crit drinkers. **(A)** IntelliCage setup and experimental timeline. Mice (n = 51) were habituated to the cage, trained to drink alcohol (4-12%) and AUD-related behaviors were tested: motivation to drink alcohol (M), extinction of alcohol seeking during withdrawal (E), alcohol seeking during cue relapse (CR), alcohol drinking during alcohol relapse (AR), and alcohol seeking during a persistence test (P). During the periods of free access to alcohol (FA) mice had unlimited access to alcohol (10%). Alcohol availability was signaled by a cue light in a reward corner. **(B)** Spearman correlations between AUD Index and AUD score and **(C)** frequency of < 2 crit and ≥ 2 crit drinkers. **(D)** Spearman correlations between AUD Index (AI) and AUD-related behaviors. Each dot on the graphs represents one animal. Linear regression lines ± 95% confidence intervals are shown; Spearman correlation (r) and ANCOVA results are given for raw data. **(E-F)** Spearman correlations (r) between AUD score (AS) and AUD-related behaviors and principal component analysis (PCA) of AUD behaviors for **(E)** < 2 crit and **(F)** ≥ 2 crit mice. Each dot on the PCA graphs represents one behavioral measure [motivation to drink alcohol (M), extinction of alcohol seeking during withdrawal (E) and cue relapse (CR), alcohol drinking during alcohol relapse (AR), and alcohol seeking during a persistence test (P)]. **(G-H)** Spearman correlations between AUD Index, **(G)** mice activity and **(H)** alcohol consumption during 4-12% and FA phases. Each dot on the graphs represents one animal. Linear regression lines ± 95% confidence intervals are shown; Spearman correlation (r) and ANCOVA results are given for raw data.

Retrospective analysis of the mice behavior showed that the ≥ 2 crit group had higher AUD score as well as scores in all AUD tests (**Supplementary Figure** 1) as compared to < 2 crit animals. AUD index correlated with AUD score (**Figure** 1C) and all AUD-related behaviors (**Figure** 1D). Moreover, when < 2 crit (n = 36) and ≥ 2 crit mice (n = 22) were analyzed separately, we found that for the ≥ 2 crit group all AUD behaviors predicted AUD score, they generally correlated with each other and heavily loaded on the main factor (PC1) in the principal components analysis (PCA) suggesting that these behaviors are measures of a single factor that may reflect compulsive alcohol use (**Figure** 1E, **Supplementary Tables** 1-2). In particular, extinction of alcohol seeking during withdrawal and persistence (Spearman r = 0.70 for both) were the best predictors of AUD-prone phenotype. On the other hand, in the < 2 crit group AUD behaviors did not correlate with AUD score, only cue relapse correlated with motivation (**Figure** 1E, left) and AUD behaviors loaded differently on principal components in PCA (**Figure** 1E, right; **Supplementary Tables** 3-4) indicating that they are driven by different factors. Hence, tight correlation between AUD behaviors characterises only AUD-prone mice, a fraction of mice drinking alcohol. Finally, despite the profound difference in addiction-like behavior scores between < 2 crit and ≥ 2 crit mice, AUD index did not predict alcohol consumption during FA or mice activity (total NPs) (**Figure** 1F-G).

Thus, the AUD model allowed for the identification of the mice that demonstrate a consistent AUD-like phenotype and AUD-resistant drinkers. As extinction of alcohol seeking during alcohol withdrawal was one of the best predictors of AUD phenotype in our model, in the following step we focused on transcriptomic differences between the < 2 crit and ≥ 2 crit mice following withdrawal.

### Differentially expressed genes in the hippocampus of ≥ 2 crit and < 2 crit mice during extinction of alcohol seeking

We hypothesized that transcriptomic differences drive the variance between the < 2 and ≥ 2 crit mice in extinction of alcohol seeking during alcohol withdrawal. To test this hypothesis 16 mice were trained to drink alcohol in the IntelliCages. For the molecular analysis ten individuals with the highest (≥ 2 crit, n = 5) and lowest AUD index (< 2 crit mice, n = 5) were selected (**Figure** 2A). They differed in AUD score as well as all AUD behaviors including extinction of alcohol seeking during withdrawal (**Figure** 2B and **Supplementary Figure** 2). The hippocampus and amygdala tissue was collected immediately after the second alcohol extinction test (day 90, **Figure** 2A), total RNA was extracted and used for a new generation high-throughput RNA sequencing (RNA-seq). We focused on these brain regions as the hippocampus has been implicated in context-induced alcohol and drug seeking during withdrawal ^26–28, 31–34^, while the amygdala, use here as a control region, was implicated in alcohol consumption despite negative consequences, alcohol choice over natural rewards, alcohol motivation as well as cue relapse, rather then alcohol seeing in alcohol-predicting contexts ^11, 16, 23, 24, 35, 45^.

**Figure 2.**
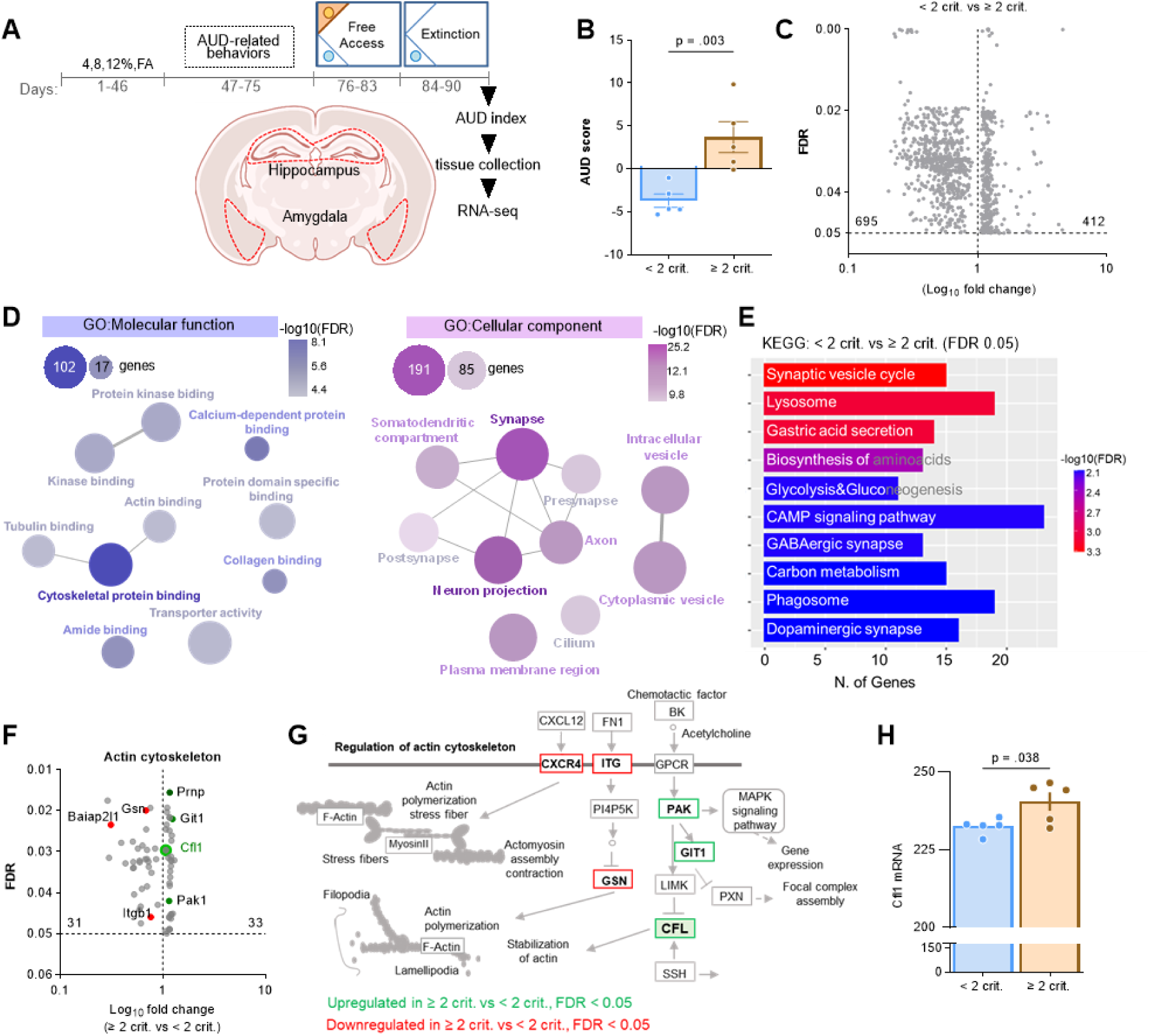
Transcripts related to cytoskeleton and synaptic function are differentially expressed in the hippocampus of the < 2 crit and ≥ 2 crit drinkers during extinction of alcohol seeking. **(A-B)** Experimental timeline. Mice were trained to drink alcohol in the IntelliCages (n = 16), classified as < 2 crit (n = 5) and ≥ 2 crit drinkers (n = 5) and sacrificed after a second alcohol extinction test (day 90) (AUD score: t(8) = 3.78, p = 0.003). Hippocampus and amygdala tissue was dissected from fresh brains for RNA-seq analysis. **(C)** A volcano plot illustrating DEGs in the hippocampus of < 2 crit and ≥ 2 crit drinkers. Only genes with FDR < 0.05 are shown. **(D)** Gene ontology analysis of molecular function (GO:MF) and cellular components (GO:CC) based on the genes deregulated (both up- and downregulated) in the hippocampus of < 2 crit vs. ≥ 2 crit drinkers (FDR cutoff 0.05). The network shows pathways in nodes. Two nodes are connected if they share 20% or more genes. Darker nodes are more significantly enriched gene sets while bigger nodes represent larger gene sets. Thicker connection between nodes represents more overlapped genes. **(E)** KEGG pathway analysis of the deregulated genes (both up- and downregulated) in the hippocampus of < 2 crit vs. ≥ 2 crit drinkers (FDR cutoff 0.05). **(F)** A volcano plot illustrating DEGs from Cytoskeletal protein binding,Tubulin binding and Actin binding nodes (GO:MF) in the hippocampus of < 2 crit vs. ≥ 2 crit drinkers. **(G)** Regulation of actin cytoskeleton pathway (KEGG) with indicated DEGs in the hippocampus of < 2 crit and ≥ 2 crit drinkers. **(H)** Cofilin (Cfl) mRNA levels in the hippocampus of < 2 crit and ≥ 2 crit drinkers (t(8)=2.49, p = 0.038).

The hippocampal transcriptome analysis of ≥ 2 crit and < 2 crit mice yielded 1107 differentially expressed genes (DEGs); 412 genes were upregulated and 695 downregulated in ≥ 2 crit as compared to < 2 crit animals (**Figure** 2C). On the other hand, we found only 4 DEGs in the amygdala; 1 gene (Snora2b) was upregulated and 3 transcripts (Prl, Gm25014 and Gm23711) downregulated in the ≥ 2 crit as compared to < 2 crit drinkers. Therefore, in the following steps of the analysis we focused on the hippocapal transcriptom.

Hippocampal DEGs were classified according to their molecular function (MF) and cellular component (CC) categories by the gene-ontology (GO) platform ^46^. GO enrichment analysis of DEGs mapped a large proportion of the genes into the *cytoskeletal function* (GO:MF, 102/987 genes, FDR = 5.10E^-12^) and *synapse* localisation (GO:CC, 191/1465 genes, FDR = 3.20E^-39^) (**Figure** 2D, **Supplementary Table** 5-6). According to the Kyoto Encyclopedia of Genes and Genomes (KEGG) ^47^ the *synaptic vesicle cycle* pathway was the most differentially expressed (**Figure** 2E, **Supplementary Table** 7) (15/77 genes, FDR = 0.00048). These results indicate that the reorganization of the cytoskeleton and changes in synaptic function in the hippocampus may contribute to differences in extinction of alcohol seeking between the < 2 and ≥ 2 crit mice.

### Cfl is upregulated in the hippocampus of the ≥ 2 crit mice during extinction of alcohol seeking

Among the top DEGs associated with the cytoskeleton function, we found upregulation of cofilin (Cfl) transcripts in the ≥ 2 crit mice as compared to < 2 crit animals (**Figure** 2F-H). Previous studies have shown that Cfl severs actin filaments, leading to increased actin cytoskeletal dynamics ^48^. This mechanism not only regulates postsynaptic function but also synaptic vesicle mobilization and exocytosis ^49–51^. Additionally, active Cfl can bind to F-actin and form stable actin rods, which can impede axonal trafficking ^52^. Since RNA-seq analysis suggests that these processes may be dysregulated in the hippocampus of ≥ 2 crit mice during alcohol withdrawal (**Figure** 2D-E) we chose to focus on Cfl in the subsequent steps of our study.

To verify distinctive expression of Cfl in the ≥ 2 crit and < 2 crit groups during extinction test, mice were trained to drink alcohol in the IntelliCages. The ≥ 2 crit and < 2 crit animals were identified and they were sacrificed after 7-day alcohol withdrawal (extinction, day 90) (**Supplementary Figure** 3). The brains were sliced and immunostained with specific antibodies. We analyzed Cfl levels on the brain slices as integrated mean gray values of the microphotographs. Significant upregulation of Cfl in the ≥ 2 crit, as compared to < 2 crit mice, was observed in the dentate gyrus of the hippocampus (DG), but not CA1 area, basolateral amygdala (BLA), central nucleus of the amygdala (CeA), nucleus accumbens (NAc) and caudate putamen (CaPu) (**Supplementary Figure** 3D-E).

To confirm this observation the experiment was repeated with a new cohort of mice. The ≥ 2 crit and < 2 crit mice were sacrificed during free alcohol access period (alcohol, day 83) or after 7-day extinction test (extinction, day 90) (**Supplementary Figure** 4, **Figure** 3A and B). We also used alcohol-naive mice as a control. We focused on the analysis of Cfl in the DG layers: the granule cell layer (GCL), polymorphic layer of dDG (PoDG), as well as the molecular layer of GC dendrites (ML) (**Figure** 3C). Overall, Cfl levels were increased in all mice drinking alcohol as compared to alcohol-naive animals. Furthermore, the levels of Cfl were increased in the ML and PoDG after extinction test in the ≥ 2 crit mice, as compared to the < 2 crit extinction animals and the ≥ 2 crit alcohol group (**Figure** 3D).

**Figure 3.**
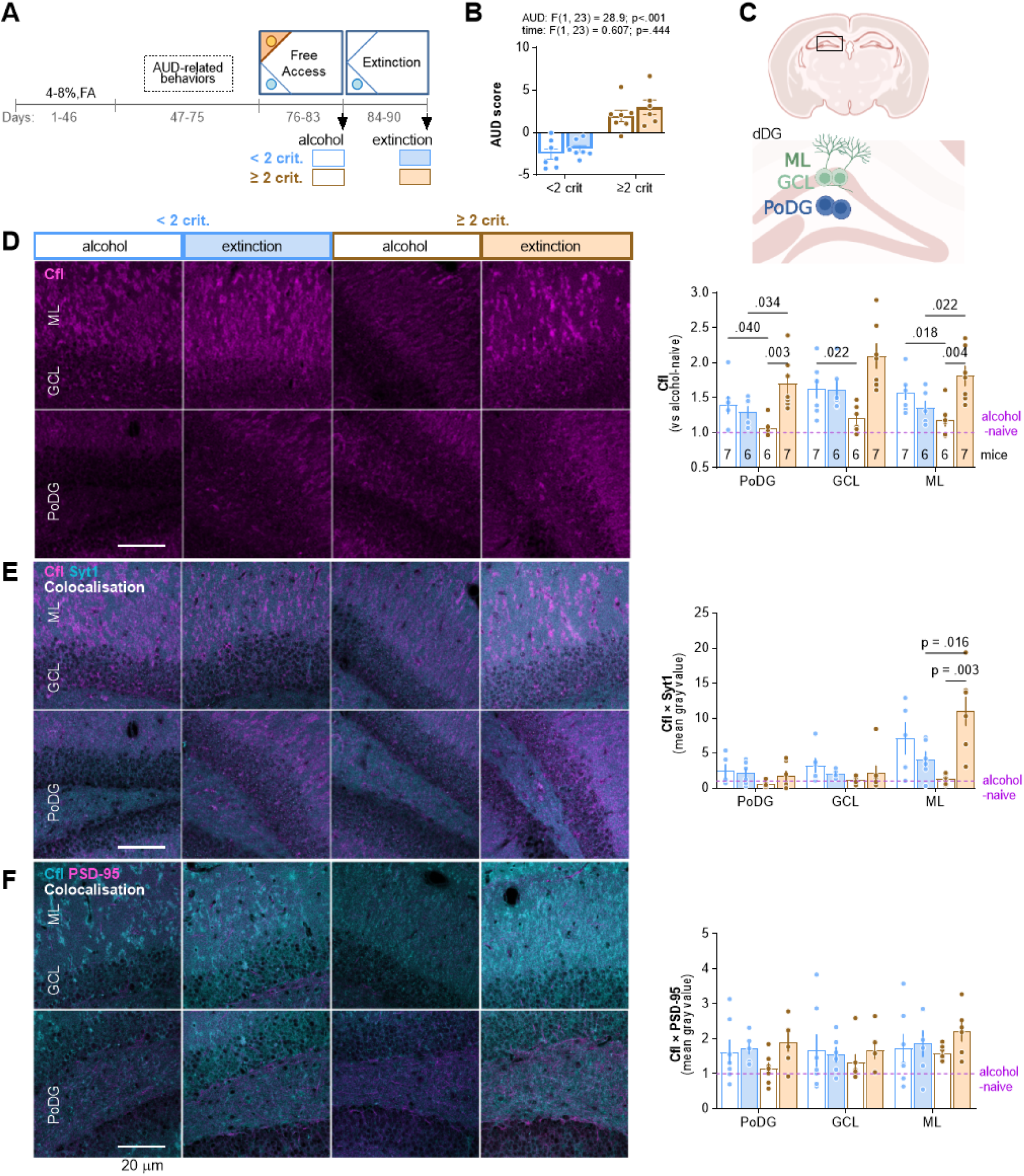
Cfl is upregulated in DG of the ≥ 2 crit animals during extinction of alcohol seeking. **(A-B)** Experimental timeline. Mice were trained to drink alcohol and AUD-related behaviors were tested. **(B)** AUD scores were calculated to identify the < 2 crit vs ≥ 2 crit drinkers. < 2 crit vs. ≥ 2 crit mice significantly differed in alcohol seeking during withdrawal (Mann Whitney test, U=2). Mice were sacrificed and perfused after free alcohol access period (day 83) or alcohol withdrawal (day 90). The brains were cut and brain sections immunostained to detect cofilin (Cfl). **(C)** Schematic representation of the analyzed dDG regions. GCL, granule cell layer; PoDG, polymorphic layer of DG; ML, molecular layer. **(D)** The analysis of Cfl fluorescent immunostaining. Representative microphotographs and summary of data (repeated measures three-way ANOVA with Šídák’s multiple comparisons test, effect of region: F(1,40, 30,8) = 16.6, p < 0.001; effect of phenotype: F(1, 23) = 0.169, p = 0.685; effect of time: F(1, 23) = 7.33, p = 0.013; effect of region × AUD: F(2, 44) = 0.0189, p = 0.981; effect of region × time: F(2, 44) = 3.36, p = 0.044; effect of phenotype × time: F(1, 23) = 14.2, p = 0.001. **(E)** The analysis of Cfl colocalization with pre-synaptic marker, synaptotagmin 1 (Syt1). Representative microphotographs and summary of data (repeated measures three-way ANOVA with Šídák’s multiple comparisons test, effect of region: F(1.71, 34.3) = 26.3, p < 0.001; effect of phenotype: F(1, 22) = 0.598, p = 0.447; effect of time: F(1, 22) = 1.88, p = 0.184; effect of region × phenotype: F(2, 40) = 0.919, p = 0.184; effect of region × time: F(2, 40) = 4.04, p = 0.025; effect of phenotype × time: F(1, 22) = 11.8, p = 0.002. **(F)** The analysis of Cfl colocalization with a postsynaptic marker, PSD-95. Representative microphotographs and summary of data (repeated measures three-way ANOVA, effect of region: F(1.84, 40.5) = 4.59, p = 0.018; effect of phenotype: F(1, 22) = 0.0296, p = 0.865; effect of time: F(1, 22) = 1.34, p = 0.259; effect of region × phenotype F(2, 44) = 0.836, p = 0.440; effect of region × time: F(2, 44) = 1.27, p = 0.292; effect of phenotype × time: F(1, 22) = 1.06, p = 0.315. Mean ± SEM are shown. For B, D-F each dot on the graphs represents one animal.

As RNA-seq analysis suggested deregulation of the synaptic proteins and proteins related to synaptic vesicle cycle in the ≥ 2 mice (**Figure** 2E), we also analyzed colocalization of Cfl with synaptotagmin 1 (Syt1) (Ca^2+^ sensor in the membrane of the pre-synaptic axon terminal involved in both synaptic vesicle docking and fusion with the presynaptic membrane; upregulated in RNA-seq data, **Supplementary Table** 1) and PSD-95/Dlg4 (post-synaptic scaffold protein; upregulated in RNA-seq data, **Supplementary Table** 1) (**Figure** 3E-F). Overall, synaptic Cfl levels (Cfl colocalizing with Syt1 and PSD-95) were increased in mice drinking alcohol as compared to alcohol-naive mice. We also observed increased levels of Cfl colocalized with Syt1 in the ML in the ≥ 2 crit mice after extinction, as compared to the ≥ 2 crit alcohol group and the < 2 crit extinction animals (**Figure** 3E). There was no significant effect of the training and AUD on the levels of Cfl co-localised with PSD-95 (**Figure** 3F). Altogether, our analysis shows that alcohol training upregulates Cfl in DG. Furthermore, extinction of alcohol seeking upregulates Cfl in PoDG and ML in the ≥ 2 crit mice as compared to the < 2 group; and the upregulated Cfl in ML colocalized with the pre-rather than post-synaptic compartments.

### Extinction of alcohol seeking impairs ML synaptic function in ≥ 2 crit mice

To test whether extinction of alcohol seeking induces synaptic changes in the ML of the ≥ 2 crit mice, we trained a new cohort of animals to drink alcohol in the IntelliCages. As previously, ≥ 2 and < 2 crit mice were identified and sacrificed during FA (alcohol) or after extinction test (**Figure** 4A and B, **Supplementary Figure** 5). Alcohol-naive mice were used as a control. Field excitatory postsynaptic potentials (fEPSPs) were recorded to evaluate synaptic function by measuring input-output and paired-pulse ratio (PPR) in the ML synapses of acute hippocampal slices when axons terminating in the ML were stimulated by monotonically increasing stimuli (**Figure** 4C).

**Figure 4.**
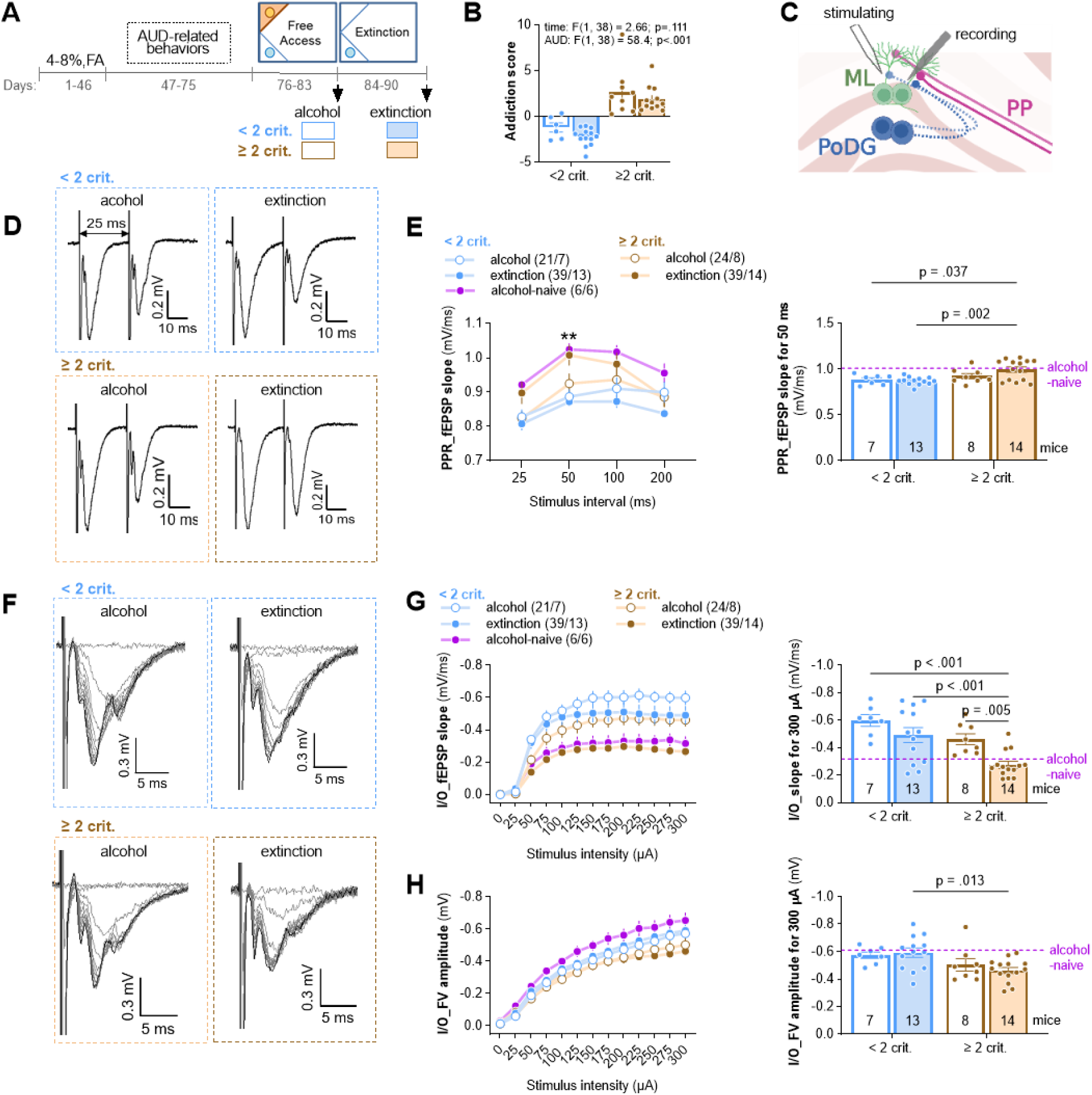
Extinction of alcohol seeking impairs synaptic transmission in the ML of the ≥ 2 crit mice. **(A-B)** Experimental timeline. Mice were trained to drink alcohol and **(B)** AUD scores were calculated to identify the < 2 crit vs ≥ 2 crit drinkers. < 2 crit vs. ≥ 2 crit mice significantly differed in alcohol seeking during withdrawal (Mann Whitney test, U=2). Mice were sacrificed after the period of free access to alcohol (day 83) or alcohol withdrawal (day 90). Alcohol-naive mice trained in the IntelliCage were used as a control. **(C)** Schematic representation of dDG. Field excitatory postsynaptic potentials (fEPSPs) were recorded in the molecular layer of dDG (ML_dDG) in response to the stimulation of the axons terminating in the ML: perforant pathway (PP) axons from the entorhinal cortex and axons from the contralateral polymorphic layer of DG (PoDG). **(D-H)** The analysis of synaptic responses. **(D)** Example of PPR traces of fEPSPs with 25 ms interstimulus interval. **(E, right)** Summary of data for PPR fEPSP slope recorded for different inter-stimulus intervals (25, 50, 100, 200 ms) (top, repeated measure two-way ANOVA, effect of stimulation intensity, F(2.29, 86.9) = 21.7, p < 0.001; treatment effect, F(3, 38) = 2.94, p = 0.046; treatment × stimulation interaction, F(9, 114) = 2.50, p = 0.012). **(E, left)** Summary of data for PPR fEPSP slope recorded for 50 ms interval (top, two-way ANOVA with Šídák’s multiple comparisons test, phenotype effect, F(1, 38) = 9.56, p = 0.004, time effect, F(1, 38) = 1.13, p = 0.295). **(G)** Representative fEPSPs traces evoked by stimuli of different intensities. **(H, left)** Summary of data for input–output plots of fEPSP slope recorded in response to increasing intensities of stimulation (repeated measure three-way ANOVA, effect of phenotype: F(1, 37) = 18.2; p<.001; effect of time, F(1, 37) = 8.47; p = 0.006). **(H, right)** Summary of data for fEPSP slope elicited by 300 µA stimulus intensity (two-way ANOVA with Šídák’s multiple comparisons test, effect of phenotype: F(1, 38) = 15.1, p < 0.001; effect of time: F(1, 38) = 10.5, p = 0.003). **(I, left)** Summary of data for fiber volley (FV) amplitude recorded in response to increasing intensities of stimulation (repeated measure three-way ANOVA, phenotype: F(1, 38) = 6.22, p = 0.017; time: F(1, 38) = 0.162, p = 0.689). **(right)** Summary of data for FV slope elicited by 300 µA stimulus intensity (two-way ANOVA with Šídák’s multiple comparisons test, effect of phenotype: F(1, 38) = 8.84, p = 0.005; effect of time, F(1, 38) = 0.102; p = 0.751). Numbers of slices/animals per group are indicated in the legends. Data are presented as means of a group +/- SEM.

The PPR slope was significantly decreased in the alcohol mice as compared to alcohol-naive animals, and increased in the ≥ 2 crit mice sacrificed after extinction test, as compared to the < 2 crit extinction animals and the ≥ 2 crit alcohol mice (**Figure** 4D-E). This data indicates higher presynaptic release probability in the alcohol mice, as compared to the alcohol-naives, and lower in the ≥ 2 crit mice after extinction as compared to other alcohol groups. Moreover, the input-output curves for the slope of fEPSP were increased in the alcohol mice as compared to alcohol-naives, and decreased in the ≥ 2 crit extinction mice compared to the ≥ 2 crit alcohol group and < 2 crit extinction mice (**Figure** 4G-I), suggesting more synaptic transmission in the alcohol groups as compared to alcohol-naive animals, and less synaptic transmission in the ≥ 2 crit extinction mice as compared to other alcohol groups. We also observed lower fiber volley (FV) responses in the ≥ 2 crit mice as compared to the alcohol-naive and < 2 crit animals indicating less activated axons in the ≥ 2 crit mice.

Overall, higher PPR and lower input-output in the alcohol-naive mice as compared to alcohol-trained animals indicate increased synaptic function after alcohol training. However, higher PPR and lower input-output in the ≥ 2 crit mice after extinction test, as compared to the < 2 crit extinction group and the ≥ 2 crit alcohol animals, indicate weakening of the ML synapses of ≥ 2 crit mice during alcohol withdrawal. This process is likely driven by pre-synaptic changes.

### Overexpression of cofilin in PoDG weakens contralateral synapses in the ML of DG

The main inputs to the ML originate from the entorhinal cortex and contralateral PoDG (**Figure** 4C). As we observed increased Cfl levels in PoDG of the ≥ 2 crit mice after extinction (**Figure** 3D) we hypothesized that weakened synaptic transmission in the ML of the ≥ 2 crit mice after extinction test, as compared to the ≥ 2 crit mice before extinction, is driven by the increase of pre-synaptic Cfl levels in PoDG. To address this hypothesis, alcohol-naive mice were unilaterally injected into DG with adeno-associated viral vectors (AAV_2.1_) expressing cofilin with hemagglutinin tag (HA) (Cfl) under CaMKII promoter ^53^. This resulted in Cfl overexpression in the PoDG cells (Cfl_PoDG) ipsilaterally to the injection, and in the PoDG axons in the ML (Cfl_ML) contralaterally to the injection (**Figure** 5A-D). The AAV_2.1_ encoding eGFP under CaMKII promoter was used as a control. The fEPSPs were recorded to measure input-output and PPR in the ML while axons terminating in the ML were stimulated (**Figure** 5D).

**Figure 5.**
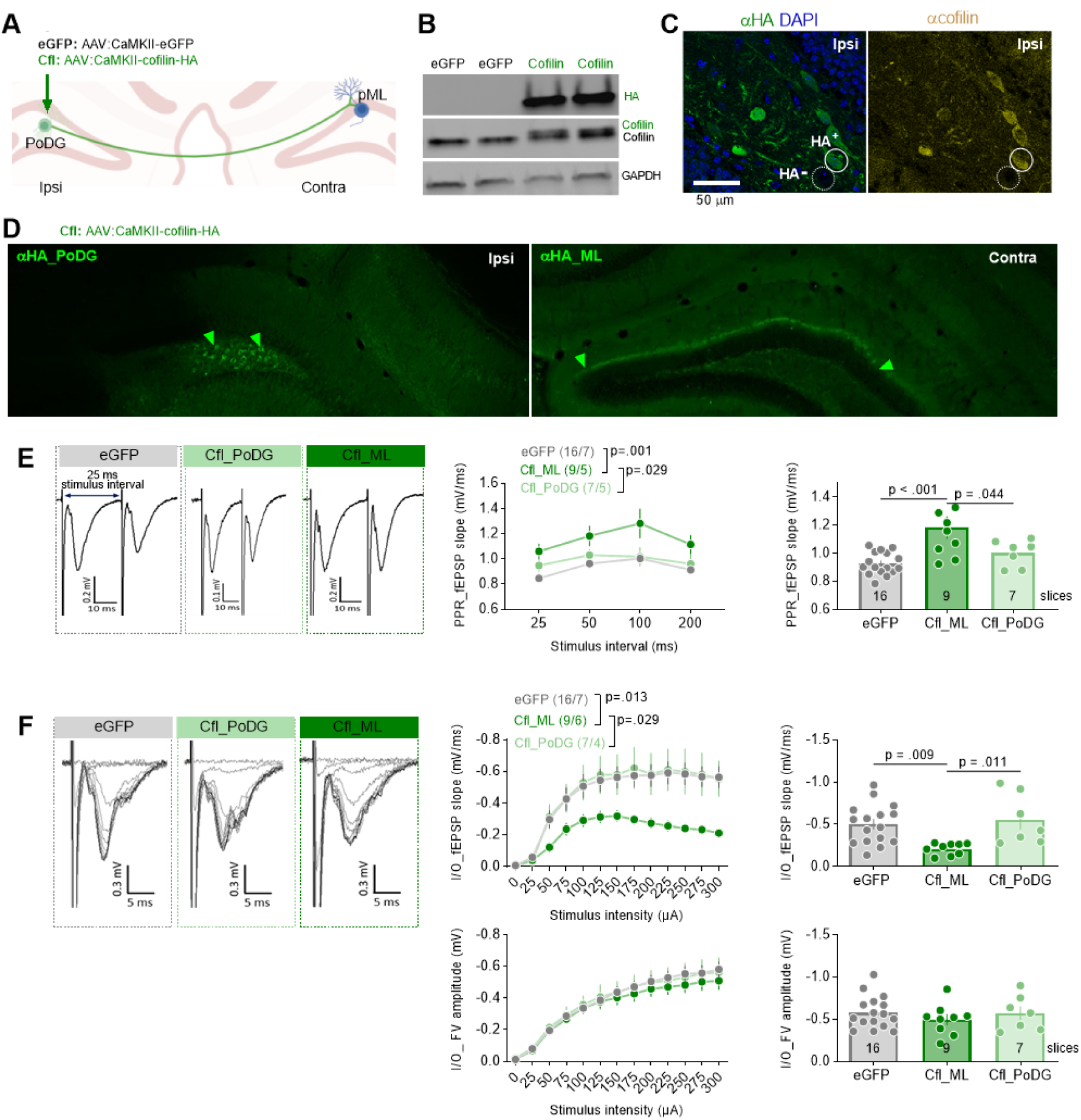
Overexpression of cofilin in PoDG weakens strength of the contralateral PoDG→ML synapses. **(A)** Mice received dDG-targeted unilateral stereotactic injections of AAV encoding cofilin-HA under CaMKII promoter (Cfl), or eGFP as a control. **(B)** Blot shows the expression of endogenous cofilin in dDG (recognized by α-cofilin antibody), and exogenous cofilin-HA protein (green, detected by α-HA antibody and as a band shifted upwards on the α-cofilin blot). GAPDH was used as a loading control for WB. **(C)** Representative microphotographs of the α-HA and α-cofilin fluorescent immunostaining in PoDG (ipsilateral to virus injection). HA-positive (HA^+^) and HA-negative (HA^-^) cells are indicated. **(D)** Representative microphotographs of α-HA fluorescent immunostaining ipsilateral to virus injection (in PoDG) and contralateral to injection (in ML). **(E-F)** The analysis of synaptic responses in ML of the brain slices with eGFP and Cfl (in PoDG or ML). Field excitatory postsynaptic potentials (fEPSPs) were recorded in ML in response to the stimulation of the axons terminating in ML (see Fig. 4C). **(E)** Example of PPR traces of fEPSPs with 25 ms inter-stimulus interval and summary of data for PPR fEPSP slope recorded for different inter-stimulus intervals (25, 50, 100, 200 ms) (repeated measure ANOVA, effect of stimulation intensity, F(2, 123) = 20.9, p < 0.001; treatment effect, F(3, 123) = 4.22, p = 0.007; treatment × stimulation interaction, F(6, 123) = 0.306, p = 0.933). **(E, right)** Summary of data for PPR fEPSP slope recorded for 50 ms interval (one-way ANOVA with Tukey’s multiple comparisons test, F(2, 29) = 9.33, p < 0.001). **(F)** Representative fEPSPs traces evoked by stimuli of different intensities and summary of data for input–output plots of fEPSP slope recorded in response to increasing intensities of stimulation (repeated measure ANOVA, effect of stimulation intensity, F(12, 372) = 63.8, p <.001; treatment effect, F(2, 31) = 2.91, p = 0.069; treatment × stimulation interaction, F(24, 372) = 2.92, p < 0.001). **(F, top right)** Summary of data for fEPSP slope elicited by 300 µA stimulus (one-way ANOVA with Tukey’s multiple comparisons test, F(2, 30) = 6.16, p = 0.006). **(F, bottom)** Summary of data for fiber volley (FV) amplitude recorded in response to increasing intensities of stimulation (repeated measure three-way ANOVA, AUD: F(1, 38) = 6.22, p = 0.017; time: F(1, 38) = 0.162, p = 0.689). **(F, bottom right)** Summary of data for FV amplitude elicited by 300 µA stimulus intensity (repeated measure ANOVA, effect of stimulation intensity, F(1,33, 38,6) = 149, p <.001; treatment effect, F(2, 29) = 0.258, p = 0.774 treatment × stimulation interaction, F(24, 348) = 0.408, p = 0.995). Data are presented as means +/- SEM. The numbers of analyzed slices and animals are indicated in the legends.

The PPR, analyzed as slope of fEPSP, was significantly increased in the Cfl_ML slices, as compared to the eGFP and the Cfl_PoDG sections (**Figure** 5E). The fEPSP slope of the input-output test was significantly decreased in the Cfl_ML slices compared to the eGFP and Cfl_PoDG slices. We did not observe the difference in the ML synaptic strength in the Cfl_PoDG slices compared to the eGFP sections (**Figure** 5F). We also did not observe any effect of the virus on FV amplitude (**Figure** 5F). Thus, overexpression of Cfl in the PoDG cells decreased the probability of synaptic release and synaptic transmission in the contralateral PoDG**→**ML synapses. This indicates that increased expression of Cfl in the PoDG**→**ML synapses is a plausible mechanism that decreases ML synaptic function in the ≥ 2 crit mice during alcohol withdrawal.

### Overexpression of cofilin in PoDG impairs extinction of alcohol seeking and increases alcohol motivation

To test whether PoDG Cfl affects AUD-related behaviors, mice were bilaterally injected with Cfl (n=13) or the control eGFP virus (n=12). Two weeks after the surgery the animals started long-term alcohol training in the IntelliCages (**Figure** 6A). Post-training analysis of the hippocampal sections showed that PoDG cells expressing Cfl had higher levels of Cfl and F-actin as compared to the non-transduced cells analyzed in the same animals [Cfl(-)]. Thus Cfl affected the actin cytoskeleton (**Figure** 6B-C).

**Figure 6.**
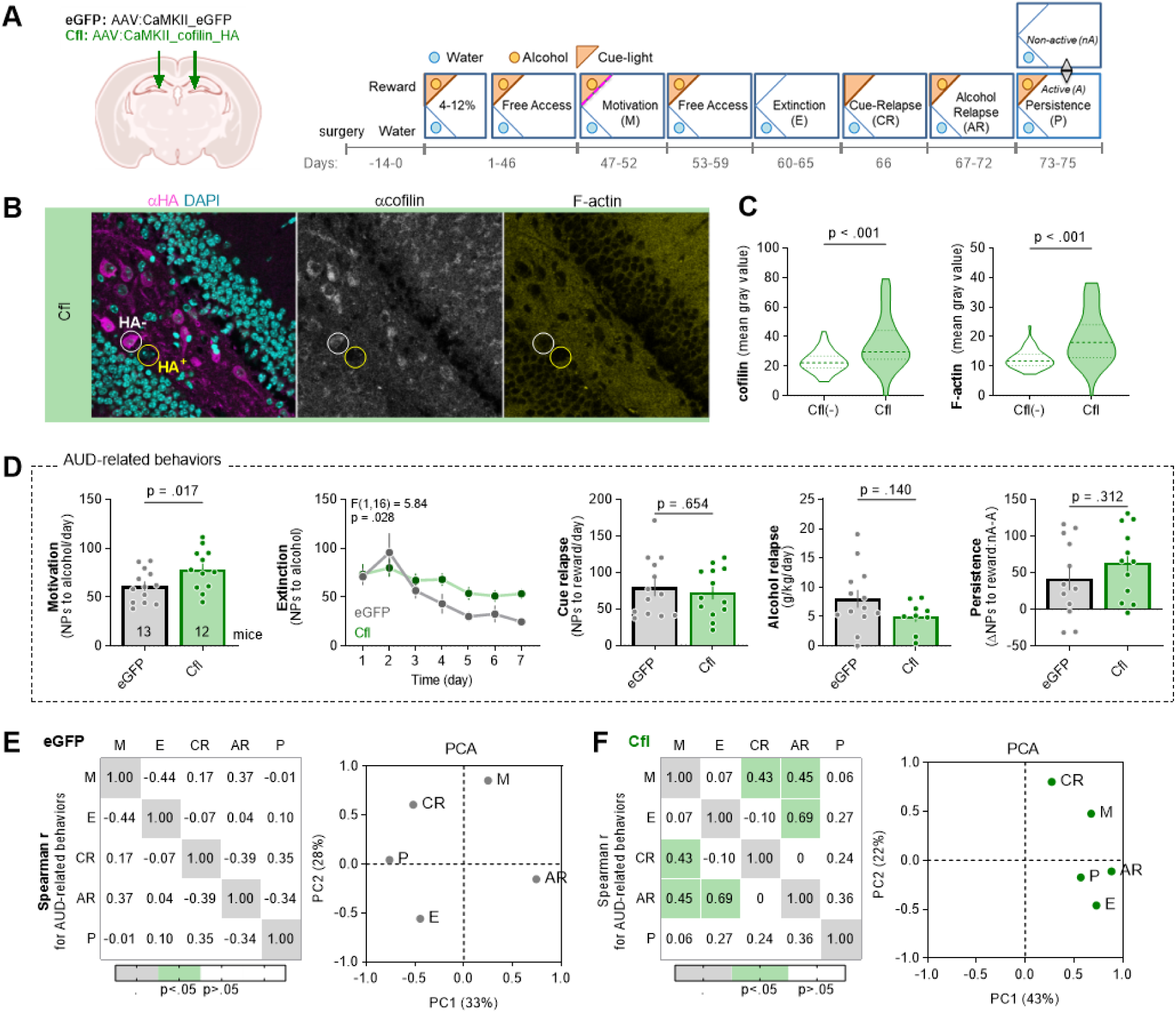
Overexpression of Cfl in PoDG impairs extinction of alcohol seeking and increases alcohol motivation. **(A)** Experimental timelines and IntelliCage setups. Mice received dDG-targeted bilateral stereotactic injections of AAV2.1 encoding cofilin-HA (n = 13), or eGFP (n = 13). Mice were trained to drink alcohol (4-12% and Free access) and AUD-related behaviors were tested: motivation to drink alcohol (M), alcohol seeking during withdrawal (W) and cue relapse (CR), alcohol drinking during alcohol relapse (AR), and alcohol seeking during a persistence test (P). **(B-C)** The analysis of cofilin and F-actin levels in PoDG cells. **(B)** Representative microphotographs of double immunolabeling with α-HA and α-cofilin, combined with DAPI (nuclear marker) and F-actin fluorescent labeling. **(C)** Summary of data showing cofilin (Mann-Whitney U = 758) and F-actin levels (Mann-Whitney U = 531) in the cells expressing cofilin, or non-transduced PoDG cells analyzed in the same animals [cofilin(-)]. Cells were detected based on DAPI staining. **(D)** Summary of data showing individual scores in AUD-related behaviors: motivation to alcohol (t(24) = 3.09), extinction of alcohol seeking during withdrawal (repeated measures one-way ANOVA, effect of virus: F(1,16) = 5.84, p = 0.028), alcohol seeking during cue relapse (t(24) = 0.455), alcohol consumption during alcohol relapse (t(23) = 1.89) and persistence in alcohol seeking (t(24) = 2.12). **(E-F)** Spearman correlations (r values) between AUD-related behaviors [motivation to drink alcohol (M), alcohol seeking during withdrawal (W) and cue relapse (CR), alcohol drinking during alcohol relapse (AR), and alcohol seeking during a persistence test (P)], and principal component analysis (PCA) of AUD behaviors for **(E)** eGFP and **(F)** Cfl mice.

Overexpression of Cfl in PoDG increased motivation for alcohol and impaired extinction of alcohol seeking during withdrawal. However, it had no effect on alcohol consumption during alcohol relapse, alcohol seeking during cue relapse and persistence in alcohol seeking (**Figure** 6D). Furthermore, PoDG Cfl had no effect on mice activity in the water corner, total activity and liquid consumption as well as alcohol seeking and consumption during 4-12% and alcohol periods (**Supplementary Figure** 6). Furthermore, we observed that the AUD behaviors of eGFP mice, as for < 2 crit animals, did not correlate with each other and loaded on different components in PCA (**Figure** 6E, **Supplementary Table** 8-9) (possibly due to the fact that in the general population there are significantly more < 2 crit than ≥ 2 crit mice). On the other hand, in the Cfl group, we observed a significant correlation between motivation and cue relapse, as well as extinction and alcohol relapse (**Figure** 6F). Moreover, persistence, alcohol relapse, extinction and motivation similarly loaded on PC1 (r = 0.57 to 0.88) (**Figure** 6F, right; **Supplementary Table** 10-11) suggesting that they measure one factor that may reflect compulsive alcohol seeking. Thus our data indicate that increased Cfl expression in PoDG resulted in specific enhancement of alcohol seeking during extinction test that possibly resulted from the increased motivation for alcohol, and increased correlation between AUD behaviours that resembled the phenotype of the ≥ 2 crit mice.

## DISCUSSION

We employed an extensive mouse model of AUD in the social context of IntelliCages to identify AUD-prone and -resistant animals - mice positive for ≥ 2 crit and < 2 crit of AUD, respectively. Next, using RNA-seq, we characterized differences between the hippocampal and amygdala transcriptomes of the ≥ 2 crit and < 2 crit animals during alcohol withdrawal. The differential expression of the hippocampal genes related to the reorganization of the actin cytoskeleton and synaptic vesicles cycle (e.g. Cfl1) significantly contributed to the distinction between the phenotypes. We also observed decreased function of the ML-DG synapses in the ≥ 2 crit drinkers during withdrawal, and such changes were not observed in the < 2 crit animals. Overexpression of Cfl in PoDG mimicked some aspects of the ≥ 2 crit phenotype including impaired function of the ML synapses, increased alcohol motivation, impaired extinction of alcohol seeking during withdrawal, and increased correlation between AUD-related behaviors.

RNA-seq enables discovery of novel molecular mechanisms of psychiatric disorders. In the context of AUD, massive transcriptomic analyses were conducted so far using either the brain tissue of AUD patients ^54–56^ or the animals with the alcohol consumption history ^57–59^. However, these approaches have important limitations. By analyzing human tissue, one cannot distinguish between the transcripts which contribute to the development of AUD and those that are altered only at the advanced stages of the disease. On the other hand, the transcriptomic analyses in animal models commonly used animals exposed to alcohol and alcohol-naive controls, without AUD diagnosis of the tested individuals. Hence, such comparisons did not allow for the distinction of the transcripts specific to alcohol exposure from those involved in AUD progression. Here, we analyzed for the first time the transcriptomic differences between the AUD-prone and AUD-resistant animals, all with free access to alcohol. These animals significantly differ in all tested AUD behaviours, but not alcohol consumption and overall activity. We found over 1000 transcripts differentially expressed in the hippocampus and only 4 in the amygdala after alcohol withdrawal. In particular, we observed differences in the hippocampal transcripts related to the cytoskeleton rearrangement, synapses, and synaptic vesicles cycle. Low number of DEGs in the amygdala during alcohol withdrawal is not surprising as this brain region was previously linked with regulation of alcohol consumption and response to alcohol-predicting cues rather than alcohol seeking in alcohol contexts ^11, 16, 23, 24, 35, 45^.

Rearrangement of the actin cytoskeleton has been linked with AUD ^60^. However, the studies in animal models associated actin binding proteins and actin cytoskeleton mostly with sensitivity to sedative effects of ethanol ^61–63^ and ethanol consumption ^64^. To our knowledge, only one study so far linked actin-binding protein, Prosapip1, with alcohol seeking and reward ^65^. Thus, the role of actin cytoskeleton in the regulation of AUD-related behaviors beyond alcohol consumption remains poorly understood. Here we found that the differential expression of the hippocampal transcripts related to actin cytoskeleton distinguished the ≥ 2 crit and < 2 crit mice during alcohol withdrawal. In particular, we show that Cfl (a key molecule regulating actin cytoskeleton and synaptic physiology ^66^) expression is increased during alcohol withdrawal in the DG of the ≥ 2 crit mice. Furthermore, we could replicate some features of the ≥ 2 crit phenotype by the local overexpression of Cfl in the PoDG. Overexpression of Cfl increased alcohol motivation, impaired extinction of alcohol seeking during withdrawal and increased correlation between AUD-related behaviors. Previously, CFL1 has been implicated in the development of neurodegenerative diseases (Alzheimer’s disease and Huntington’s disease) ^67^, neuronal migration disorders (lissencephaly, epilepsy, and schizophrenia), neural tube closure defects ^68^ and memory consolidation during sleep ^53^. Mutations in CFL1 have been associated with impaired neural crest cell migration and neural tube closure defects ^69^. Here, we extend these findings by demonstrating the role of Cfl in the regulation of the core symptoms for the AUD diagnosis.

Actin-binding molecules were linked with alcohol consumption by their effects on actin cytoskeleton stabilization. For example, local deletion of Prosapip1 (actin regulatory protein) in the nucleus accumbens results in decreased F-actin levels in the animals treated with alcohol, compared to the alcohol-treated control mice, as well as decreased alcohol consumption ^65^. On the other hand, mice lacking Eps8 show increased ethanol consumption, while Eps8 null neurons are resistant to the actin-remodeling activity of NMDA receptors and ethanol ^62^. It remains unclear how actin-binding proteins regulate AUD-related behaviors beyond alcohol consumption. Here, we observed that overexpression of Cfl in PoDG increased PPR in the contralateral ML synapses and decreased fEPSP slope in input-output test, indicating impaired function of the ML synapses that is likely driven by lower presynaptic release probability. Similar physiological changes were observed in the ≥ 2 crit mice after extinction test, as compared to the < 2 mice sacrificed at the same time point. Thus increased levels of Cfl in the ML of ≥ 2 crit mice is likely the mechanism of synaptic weakening observed in these mice during withdrawal. In its unphosphorylated, active state, cofilin severs actin filaments and increases actin cytoskeletal dynamics ^48^. By this mechanism Cfl regulates both post-synaptic function and synaptic vesicle mobilization and exocytosis ^49–51^. Moreover, active Cfl may also bind to F-actin and form stable actin rods that block axonal trafficking ^52^. Thus, the impaired function of the ML synapses after expression of Cfl in PoDG of the alcohol-naive mice, and in the ≥ 2 crit mice during alcohol withdrawal, may be driven by deregulation of synaptic vesicles exocytosis or/and generation of actin rods that impair axonal trafficking. The role of pre-synaptic compartments in the hippocampus in the regulation of extinction of alcohol seeking is also supported by our RNA-seq data showing differential expression of the genes related to axonal projections and synaptic vesicle cycle between the ≥ 2 crit and < 2 crit mice. Alternatively, Cfl may impair DG circuit by inhibiting PoDG postsynaptic compartments. However, this hypothesis is less likely as we did not observe significant changes in the levels of Cfl colocalization with PSD-95 in the ≥ 2 crit mice after extinction.

Our findings add up to previous studies showing that impaired synaptic transmission in DG drives AUD-related behaviors ^21, 22^. In particular, our former studies found that the frequency of silent synapses (that lack functional AMPA receptors) generated during cue relapse in ML positively correlates with AUD index, while downregulation of PSD-95 in the granule cells in DG drives excessive alcohol seeking during cue relapse ^21^. Here we extend these findings demonstrating the role of pre-synaptic compartments in DG in the regulation of alcohol motivation and extinction of alcohol seeking during withdrawal. Our data are in agreement with former studies showing the role of DG in drug motivation ^29, 30^ as well as extinction of memories and learning about contingencies in the context ^70–72^. Interestingly, our data also indicate that Cfl in PoDG increases overall correlation between AUD-related behaviors, the phenomenon also observed in the ≥ 2 crit vs < 2 crit mice, AUD patients vs healthy individuals, or patients with mild vs severe AUD diagnosis. In particular, by manipulating Cfl levels in PoDG we observed increased correlations between persistence, extinction and alcohol relapse. These behaviors loaded on one major component in PCA suggesting that they measure one factor that may reflect loss of control over alcohol use, and have a common neurobiological root related to the function of Cfl in DG.

Overall, by employing a multidimensional AUD model in conjunction with genomic, electrophysiological, and biochemical analyses, we have identified a novel molecular mechanism that drives increased alcohol motivation and impairs the extinction of alcohol-seeking behavior during withdrawal. This mechanism is specific to individuals prone to AUD behaviours and involves the upregulation of Cfl at DG pre-synapses, leading to a weakening of synaptic transmission during alcohol withdrawal. Collectively, these findings may pave the way for novel therapeutic strategies aimed at enhancing synaptic transmission in patients diagnosed with AUD.

## MATERIALS AND METHODS

### Subjects

Ten-week old female C57BL/6J mice were purchased from the Medical University of Bialystok, Poland. We used only females as they not only show lower levels of aggression when group-housed in the IntelliCages but more importantly drink significantly more alcohol as compared to males ^40^. Animals were housed under a 12/12 hr light/dark cycle in standard mouse home cages with *ad libitum* access to water and food. Experiments were approved by the Animal Protection Act of Poland guidelines and the 1^st^ Local Ethical Committee in Warsaw, Poland (no. 117/2016, 421/2017, 884/2019). All experiments were planned to reduce the number of animals used and to minimize their suffering.

### Animal model of AUD in the IntelliCages

After 1 week of acclimatization, the mice were injected subcutaneously (s.c.) with unique microtransponders (11.5 mm length, 2.2 mm diameter; Trovan, ID-100) under brief isoflurane anesthesia. The mice were then allowed to recover for 3 days, and the animals with properly located microtransponders were introduced to the IntelliCage system (NewBehavior AG, Zürich, Switzerland) (https://www.tse-systems.com/service/intellicage/), 15 animals per system. The IntelliCage consists of a large standard rat cage (20.5 cm high, 40 cm × 58 cm at the top, 55 cm × 37.5 cm at the base). In each corner, a triangular learning chamber is located with two bottles. To drink, only one mouse can go inside a plastic ring (outer ring: 50 mm diameter; inner ring: 30 mm diameter; 20 mm depth into outer ring) that ends with two 13 mm holes (one on the left, one on the right) that provide access to bottle nozzles. Each visit to the corner, nose-poke at the doors governing access to the bottles, and licks were recorded by the system and ascribed to a particular animal.

The training in the IntelliCage was based on published protocol ^20^ and composed of the following phases: initiation of alcohol consumption in increasing concentrations, free access to 10% alcohol, motivation test, persistence test, withdrawal, cue relapse and alcohol relapse. The timelines of the experiments are on the figures.

#### Adaptation phase

All mice had free access to all bottles with water in both active corners. All doors were open. After 24 hours, when all mice visited and licked from both corners, the doors were closed. Under a fixed ratio of reinforcement (FR 1), each nose-poke was rewarded by a 5 second access to the bottles with water.

#### Initiation of alcohol consumption

During the test, 2 corners were active, each with two bottles available. In one corner, the animals had access to water (“water corner”), and in the other (“reward corner”) animals had access to ethanol solution (Alcohol group) at increasing concentrations (4, 8, and 12% ethanol changed every 3 days, prepared from 96% ethanol and tap water) or water (Alcohol-naive mice). When alcohol (or water) was available, it was signaled by a green light turned on in the “reward corner” each time a mouse entered the corner. All liquids were available under an FR1 schedule. During free alcohol access phases mice had unlimited access to water in one corner and 4, 8 and 12% ethanol (or water) in the “reward corner” (each concentration tested for 3-5 days). Access to water and alcohol was under FR1. 10% alcohol was chosen based on maximal alcohol consumption in g/kg/day during initiation of alcohol consumption. Daily alcohol consumption (g/kg/day) was calculated with the following formula: (number of licks of 12% alcohol per day × lick volume × 0.12 × 1 g/ml) / animal weight). To calculate the average lick volume, water consumption (in μl) was measured for 3 consecutive days. The average volume of one lick was measured as the total volume consumed / number of licks. According to these calculations, an average lick volume was established as 1.94 ± 0.2 μl. In the control groups, tap water was presented in the reward corner through the whole experiment.

#### Motivation for alcohol tests

During the test, two corners were active and available to animals. The animals had to perform an increasing number of nosepokes (2, 4, 8, 12, 16, 20, 24, 28, 32, and 36) spaced by less than 1 s during one visit to open the door and be allowed for a 5 s access to the reward bottles. The number of required instrumental responses (nosepokes) increased when an animal performed 10 sets of responses of a given ratio. The tests were terminated when 90% of animals did not change the FR level during the last 24 hours. The FR level reached during the test was used as an index of motivation.

#### Extinction of alcohol seeking followed by cue and alcohol relapse

The extinction periods were signaled as the “no-reward” periods and lasted 7 days. The door to reward was closed and nosepokes to the reward corner were without scheduled consequences. Average daily number of nosepokes performed in the “reward corner” during extinction, and a difference in the average number of nosepokes during extinction vs. the last day before the test, were used as indices of alcohol seeking during withdrawal. Each extinction was followed by a 24-hour cue-relapse. A green cue light (reward-predicting cue) in the reward corner was presented each time a mouse entered the reward corner. However, nose-pokes to the reward doors had no scheduled consequences. Average daily number of nosepokes performed in the “reward corner” during cue relapse, and a difference in the average number of nosepokes during cue relapse vs. the last day of extinction, were used as indices of alcohol seeking during cue relapse. This test was followed by reward relapse when bottles with reward (alcohol or water) were added into the active “reward corner”. During the test each nose-poke into the reward door opened the door for 5 s. Amount of reward drank during the first day of relapse (number of licks) was used as an index of relapse.

#### Persistence in reward seeking tests

Each persistence test lasted 3 days, starting at the beginning of the dark phase and was composed of six, 6-hour long “active periods” (A) altered with 6-hour long “non-active periods” (nA). “Active” periods (A) were signaled by the green cue light in the reward corner. During the “active” periods, nosepokes at all doors opened the door for 5 s (FR 1). The “non-active” periods were signaled by elimination of the green cue light. During the “non-active” periods, nose-pokes on the reward side were not followed by any scheduled consequences. Number of “reward” nose-pokes performed during the test, as well as the difference of nose-pokes performed during nA and A reward periods, were used as indices of persistence.

#### Establishment of mouse subpopulations

The addiction index was calculated as previously described ^20^ and was based on five behaviors: (i) the breakpoint reached during the motivation test, (ii) persistence in alcohol seeking during the persistence test, (iii) alcohol seeking during the cue-induced relapse and (iv) extinction and (v) alcohol consumption during the alcohol relapse. An individual was arbitrarily considered positive for an AUD-like criterion when its score in the test was in the uppermost 35% of the population. The scoring allowed us to divide the mice into groups according to the number of fulfilled AUD-like criteria: “AUD-prone” who fulfilled 2 or more criteria (≥ 2 crit); “AUD-resistant” who were positive for one or none of the criteria (< 2 crit). Moreover, since the addiction index may neglect mice performance in some tests, we developed addiction score. To calculate addiction score each of the addiction-like behaviors was normalized and summed up to calculate individual addiction score according to formula: AS = (Vi (individual score) – mean(population))/SD(population). This allowed us to distinguish mice which show consistent behavioral patterns towards alcohol.

### RNA Sequencing

The hippocampal tissue was quickly dissected on ice from the fresh brain, homogenized and stored in RNAlater solution (Invitrogen, AM7020) at 4°C, for 24 hrs and kept then at −20°C till further use. Total RNA was extracted using the RNeasy Mini kit (Qiagen, 74104) according to the manufacturer’s recommendations. RNA concentration, quality and integrity were determined using a Nanodrop 1000 (Thermo Scientific) and a Bioanalyzer (Agilent). RNA libraries for sequencing were prepared using a KAPA Stranded mRNA Sample Preparation Kit (KK8401-07962169001, Kapa Biosystems, Wilmington, MA, USA). The libraries were sequenced after onboard cluster generation using HiSeq Rapid SBS Kit v2 (200 cycles) and HiSeq PE Rapid Cluster Kit v2 (Illumina) on a HiSeq 1500 (Illumina). The row data are available at GEO NCBI (GSE221166). Power (defined as the expected proportion of identified differentially expressed genes among all the truly differentially expressed genes, given at least one gene is truly differentially expressed in the data) was modeled using the ssizeRNA R package ^73^ assuming 80% of non-differential genes, significance threshold FDR = 5%, dispersion = 5% and fold change following normal distribution. Raw sequencing data were processed by trimmomatic ^74^, adapter contamination and bad quality reads or read fragments were removed. Processed reads were mapped to mm10 reference genome by STAR algorithm ^75^. Subsequently Picard MarkDuplicates algorithm was used to mark optical duplicates. Featurecounts from the subread R package ^76^ was used to count the number of fragments assigned to genes. Data normalization and statistical analysis was done by the NOIseq R package ^77^. RNA-seq quality control analysis was performed using the RSeQC package and STAR (log files). Sequencing fragments distribution, sequencing quality and percent of unique mapping fragments was evaluated. Significantly deregulated genes (FDR-adjusted p-value < 0.05) obtained from the RNA sequencing by NOISeq ^77^ analysis were used as input for the pathway analysis. By ShinyGO 0.76 ^46^ functional enrichment analysis was performed to find affected pathways in the gene ontology cellular component (GO:CC), molecular function (GO:MF) platforms and in the Kyoto Encyclopedia of Genes and Genomes database (KEGG) ^47^.

### Immunofluorescent staining

Mice were anesthetized and transcardially perfused with filtered PBS (Sigma-Aldrich) and 4% PFA (Sigma-Aldrich)/PBS. Brains were left overnight in the same fixing solution and transferred to 30% sucrose in PBS for 72 hours. Coronal brain sections (40 µm) of perfused mice (Leica CM1950 Cryostat, Leica Biosystems) were stored at −20 °C in anti-freeze buffer [PBS, 20% sucrose (Sigma–Aldrich), 15% ethylene glycol (Sigma–Aldrich), 0.05% NaN3 (Sigma–Aldrich)]. After washing from the buffer and incubation with 5% NDS (Jackson ImmunoResearch, 017-000-121) (in PBS), the slices were incubated with primary antibody (cofilin, Cell Signaling, cat. 5175S, 1:400; PSD-9, Santa Cruz, sc-6926, synaptotagmin, Novus Biologicas, NB100-1938, 1:1000) in TBS with 0.3% Triton (TBST) and 5% NDS. The next day, the sections were washed with TBST and incubated with secondary antibodies (Invitrogen, Alexa Fluor 488 #A21206, Alexa Fluor 568 #A10037, Alexa Fluor 647 #A21447). Next, the slices were washed with PBS, mounted on microscopic slides and covered with DAPI containing medium (Fluoromount-G, Invitrogen, #00-4959-52). The fluorescent staining was photographed with a confocal microscope (Zeiss LSM800, magnification 63x) by the experimenter blind to the experimental groups. For each immunostaining all pictures were taken with the same settings.

### Electrophysiology

Local field potential recording was used to analyze input-output and paired-pulse ratio (PPR) in dDG. Mice were anesthetized with Isoflurane (Iso-Vet, 1000 mg/ml) and decapitated, and the brains were rapidly dissected and transferred into ice-cold cutting artificial cerebrospinal fluid (ACSF) consisting of (in mM): 87 NaCl, 2.5 KCl, 1.25 NaH_2_PO_4_, 25 NaHCO_3_, 0.5 CaCl_2_, 7 MgSO_4_, 20 D-glucose, 75 sucrose, equilibrated with carbogen (5% CO_2/_95% O_2_). The brain was cut to two hemispheres and 350 μm thick coronal brain slices were cut using Leica VT1000S vibratome in ice-cold cutting ACSF. The resulting slices were then incubated for 15 min in cutting ACSF at 32°C, followed by minimum 60 min incubation at room temperature in recording ACSF containing (in mM): 125 NaCl, 2.5 KCl, 1.25 NaH_2_PO_4_, 25 NaHCO_3_, 2.5 CaCl_2_, 1.5 MgSO_4_, 20 D-glucose, equilibrated with carbogen.

Extracellular field potential recordings were conducted in a submerged chamber perfused with recording ACSF in RT. The synaptic potentials were evoked with a custom built stimulus isolator using a concentric bipolar electrode (FHC, 30200) placed in the perforant path. The stimulating pulses were delivered at 0.033 Hz and the pulse duration was 0.3 ms. Recording electrodes (resistance 1-4 MΩ) were pulled from borosilicate glass (WPI, 1B120F-4) with a micropipette puller (NARISHIGE, PP-830) and filled with recording ACSF. The recording electrodes were placed in the ML of dDG. Recordings were acquired with MultiClamp 700B amplifier (Molecular Devices, California, USA), digitized with Digidata 1550B (Molecular Devices, California, USA) and recorded with Clampex 10.7 software (Molecular Devices, California, USA). Input/output curves were obtained by increasing stimulation intensity by 25 μA in the range of 0-300 μA. To measure the PPR ^78^, two electric stimuli triggering presynaptic action potentials were paired with increasing interstimulus intervals (25, 50, 100, 200 ms) and simultaneously fEPSPs were recorded for each interstimulus interval. The amplitude and slope of fEPSPs were measured using AxoGraph 1.7.4 software (Axon Instruments, U.S.A).

### Stereotaxic surgery

Mice were anaesthetized with isoflurane (5% for induction, 1.5-2.0% for maintenance of general anesthesia), fixed in the stereotactic frame (51503, Stoelting, Wood Dale, IL, USA), and their body temperature was maintained using a heating pad. Stereotactic injections were performed bilaterally into the dDG region of the hippocampus using coordinates from the Bregma: ML, ±1.0 mm; AP, −2.0 mm; DV, −2.0 mm according to (Paxinos and Franklin, 2001). 0.5 µl of virus solution was microinjected through a beveled 26 gauge metal needle, attached to a 10 µl microsyringe (SGE010RNS, WPI, USA) connected to a microsyringe pump (UMP3, WPI, Sarasota, USA) and its controller (Micro4, WPI, Sarasota, USA), at a rate 0.1 µl/min. The needle was left in place for an additional 10 minutes following injection to prevent leakage of the vector. Mice were injected with adeno-associated viral vectors (AAV_2.1_) encoding wild-type cofilin protein under CaMKII promoter fused with HA (AAV2.1:CaMKII_cofilin_HA) (Cfl) (viral titer: 7.29 × 10^8^ gc/µl), or a control eGFP-coding vector (AAV2.1_CaMKII_eGFP) (eGFP) (viral titer: 6.9 × 10^8^ gc/µl).The viruses were prepared by the Laboratory of Animal Models at Nencki Institute of Experimental Biology, Polish Academy of Sciences. After the surgery, animals were given 14 days to recover before training in the IntelliCages. After training, the animals were perfused with 4% PFA in PBS and brain sections from the dorsal hippocampus were immunostained to detect Cfl and HA, and imaged with Zeiss Spinning Disc confocal microscope (magnification: 10x) to assess the extent of the viral expression and proteins level (ImageJ software).

### Western blot

Mice were decapitated under isoflurane anesthesia, hippocampi were isolated and sliced into 1 mm-thick slices. dDG was cut from the slices with a razor blade. The tissue was homogenized in ice-cold lysis buffer (25mM HEPES, pH 7.4; 500mM NaCl; 2mM EDTA; 20mM NaF; 1× protease inhibitor cocktail tablet; 0.1% (v/v) Nonidet P-40). After sonication and spinning (20000xg 4°C) the supernatant was stored at −80 °C until further analysis. For Western-blot equal amounts of total protein from each sample were mixed with a Laemmli buffer containing DTT (50 mM) and left to denature at 70°C for 10 minutes. The mixture was loaded on TGX precast gel wells, that contain trihalo compounds allowing stain-free visualization of total proteins (Bio-Rad #4568083), and ran until the loading buffer reached the bottom of the gel. The analyzed protein levels were normalized to the total protein levels. Next, the proteins were transferred to membranes. The membranes were blocked by 5% or 10% (depending on the antibody) milk diluted in TBST (Tris-buffered saline with Tween 20), and incubated with the primary antibody (cofilin, Cell Signaling, cat. 5175S, 1:3000; HA-Tag, Santa Cruz, sc-7392, 1:1000; GAPDH, Merck Millipor, No. AB2302, 1:2000) for 12 hours. After washing in TBST the membranes were incubated in a secondary antibody with HRP (anti-rabbit, Vector pI-1000, 1:5000; anti-mouse, Santa Cruz, sc-2005, 1:5000) and washed again. The membranes were visualized by G-Box apparatus using chemiluminescent reagent (Advansta, #K-12042-D10). TGX stain-free gels from Bio-Rad (#4568095) were used to standardize protein loading on gels ^79^.

### Statistical analysis

The sample sizes of the experimental groups, and details of the statistics, are placed on the graphs or in the legends. The data with normal distribution and equal variance are presented as the mean with SEM and were analyzed with Student’s t-test, one-way, two-way or repeated measure two-way ANOVA. P*ost hoc* Šídák’s and Tukey’s tests for multiple comparisons were used. Alternatively, multiple repeated Mann-Whitney’s tests corrected with Holm-Sidak method for multiple comparisons were used. Correlations were analyzed using Pearson correlation. The difference between the experimental groups was considered significant at p < 0.05. All statistical analyses were performed using GraphPad Prism 9.3.1 Software.

## Acknowledgments

This work has been supported by the European Union’s Horizon 2020 research and innovation programme under the Marie Sklodowska-Curie grant agreement no 665735 (Bio4Med) and by the funding from Polish Ministry of Science and Higher Education within 2016-2020 funds for the implementation of international projects (agreement no 3548/H2020/COFUND/2016/2) and National Science Centre (Poland) Harmonia and Opus grants (2016/22/M/NZ4/00674 and 2015/19/B/NZ4/03163) to KR. TA is supported by the Roy J. Carver Chair of Neuroscience and NIH R01 MH 087463.

## Author contributions

Conceptualization: TA, RH, KR

Methodology: RP, AS, BW, BG.

Investigation: RP, AS, ES, MP, AC, ZH.

Visualization: RP, AS, KR.

Supervision: KR

Writing—original draft: RP, AS, KR

Writing—review & editing: all authors.

## SUPPLEMENTARY FIGURES

**Supplementary figure 1.**
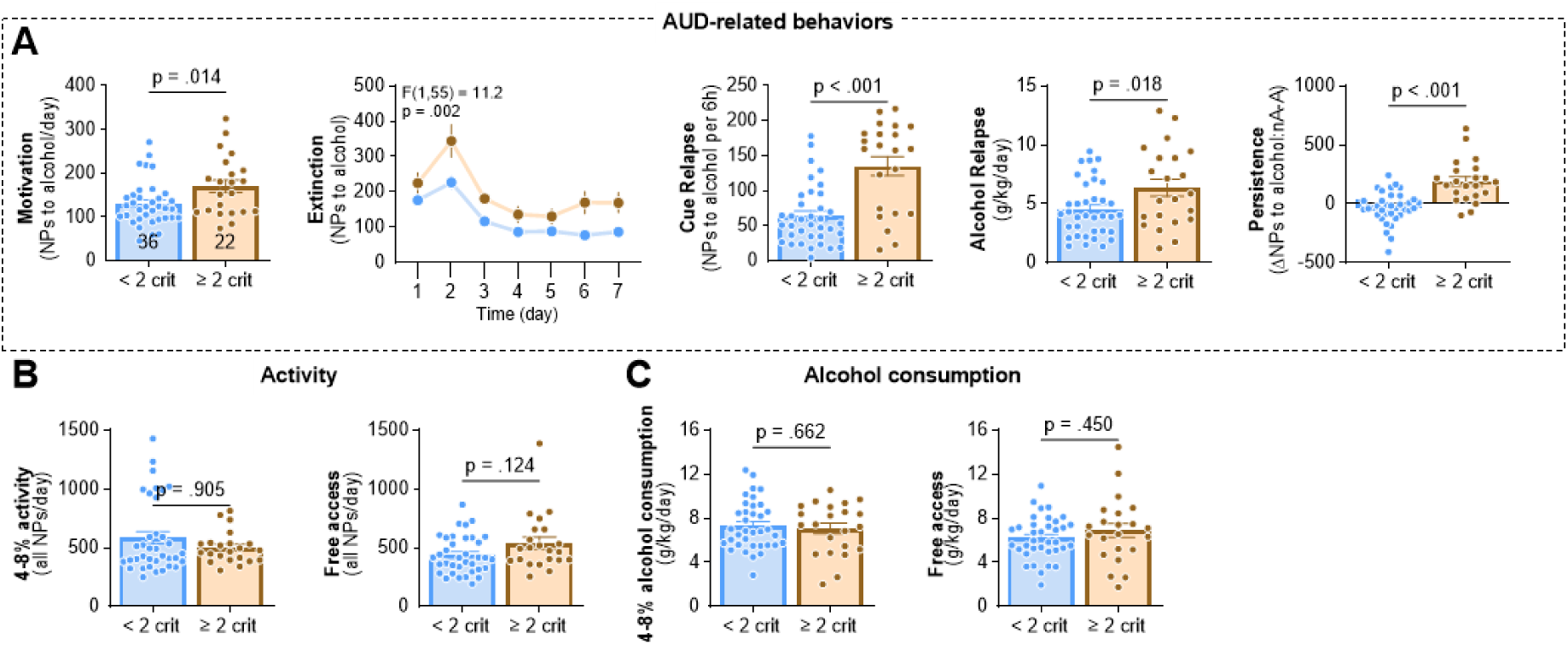
Behavioral characteristics of the < 2 crit and ≥ 2 crit mice. Mice were trained to drink alcohol (4% and 8%) and AUD-related behaviors were tested: motivation to drink alcohol (Motivation), extinction of alcohol seeking during withdrawal (Extinction) and alcohol seeking during cue relapse (Cue relapse), alcohol drinking during alcohol relapse (Alcohol relapse), and alcohol seeking during a persistence test (Persistence). During periods of free access to alcohol (FA) mice had unlimited access to alcohol (8%). ≥ 2 crit and < 2 crit mice were identified based on AUD index (AI). **(A)** Assessment of AUD-related behaviors in ≥ 2 crit and < 2 crit mice: motivation towards alcohol (t(7) = 2.62, p = 0.017), alcohol seeking during cue relapse (CR) compared to the last day of withdrawal (−1) (repeated measure ANOVA, time x AI interaction: F(1, 14) = 6.82, p = 0.021), alcohol consumption during alcohol relapse (AR) compared to the last day before withdrawal (−1) (repeated measure ANOVA, effect of time: F(1, 14) = 13.9, p = 0.002; effect of AI: F (1, 14) = 26.9, p < 0.001), persistence in alcohol seeking (t(7) = 2.28, p = 0.028) and alcohol seeking during withdrawal (repeated measure ANOVA, effect of time: F(1.19, 8.31) = 3.68, p = 0.086; effect of AI: F(1, 7) = 6.73, p = 0.036). **(B-C)** Assessment of general activity and alcohol consumption in ≥ 2 crit and < 2 crit mice.

**Supplementary figure 2.**
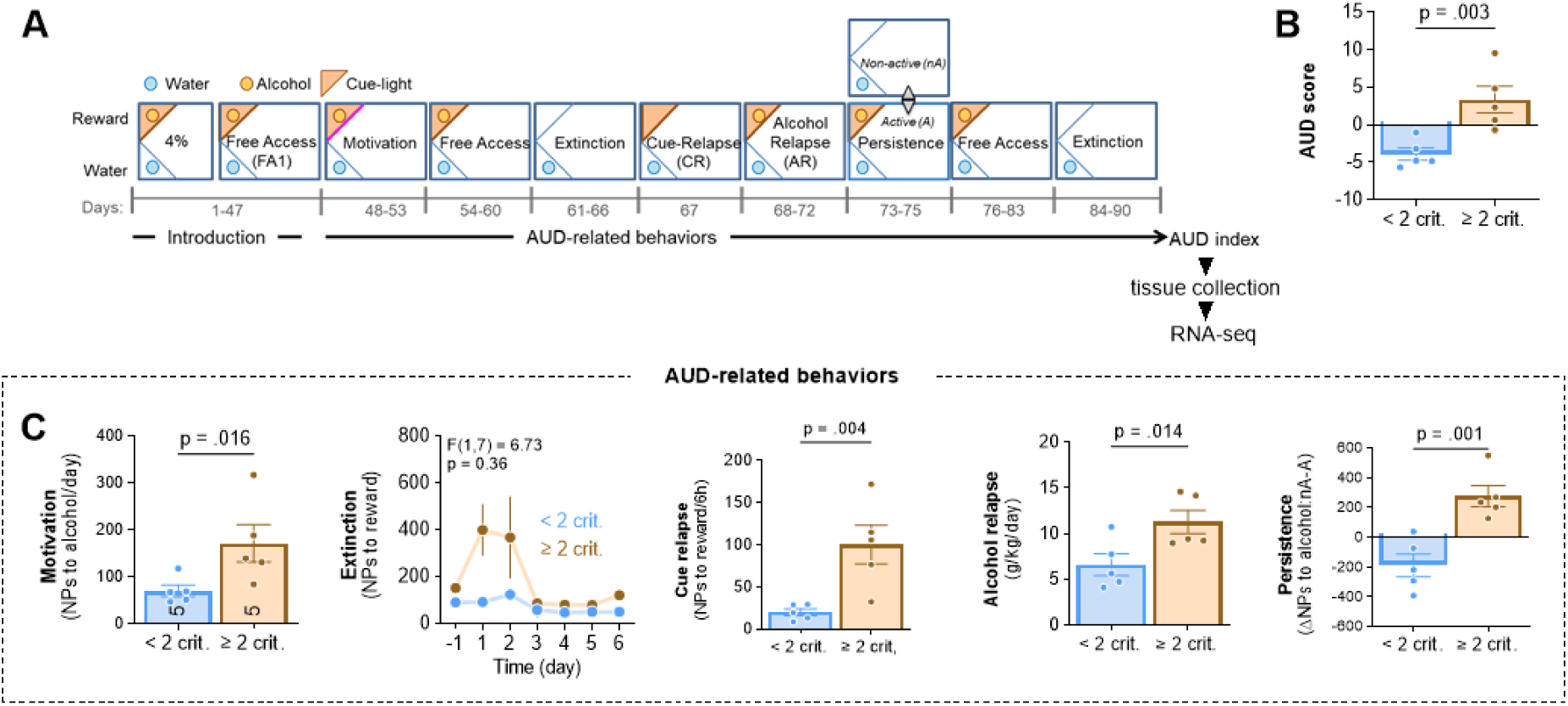
Behavioral characteristics of the < 2 crit and ≥ 2 crit mice (RNA-seq experiment). **(A)** IntelliCage setup and experimental timeline. Mice were trained to drink alcohol (4% and 8%) and AUD-related behaviors were tested: motivation to drink alcohol (Motivation), alcohol seeking during withdrawal (Withdrawal) and cue relapse (Cue relapse), alcohol drinking during alcohol relapse (Alcohol relapse), and alcohol seeking during a persistence test (Persistence). During periods of free access to alcohol (FA) mice had unlimited access to alcohol (8%). ≥ 2 crit and < 2 crit mice were identified based on AUD index (AI). **(B)** AUD score was calculated as a sum of normalized scores in AUD tests (t(8) = 3.78, p = 0.003). **(C)** Assessment of AUD-related behaviors in ≥ 2 crit and < 2 crit mice: motivation towards alcohol (Mann-Whitney U = 1), extinction of alcohol seeking during withdrawal (repeated measure ANOVA, effect of phenotype: F(1, 7) = 6.73, p = 0.036), alcohol seeking during cue relapse (CR) (Mann-Whitney U = 0), alcohol consumption during alcohol relapse (AR) (t(8) = 2.68), and persistence in alcohol seeking (t(8) = 4.41).

**Supplementary figure 3.**
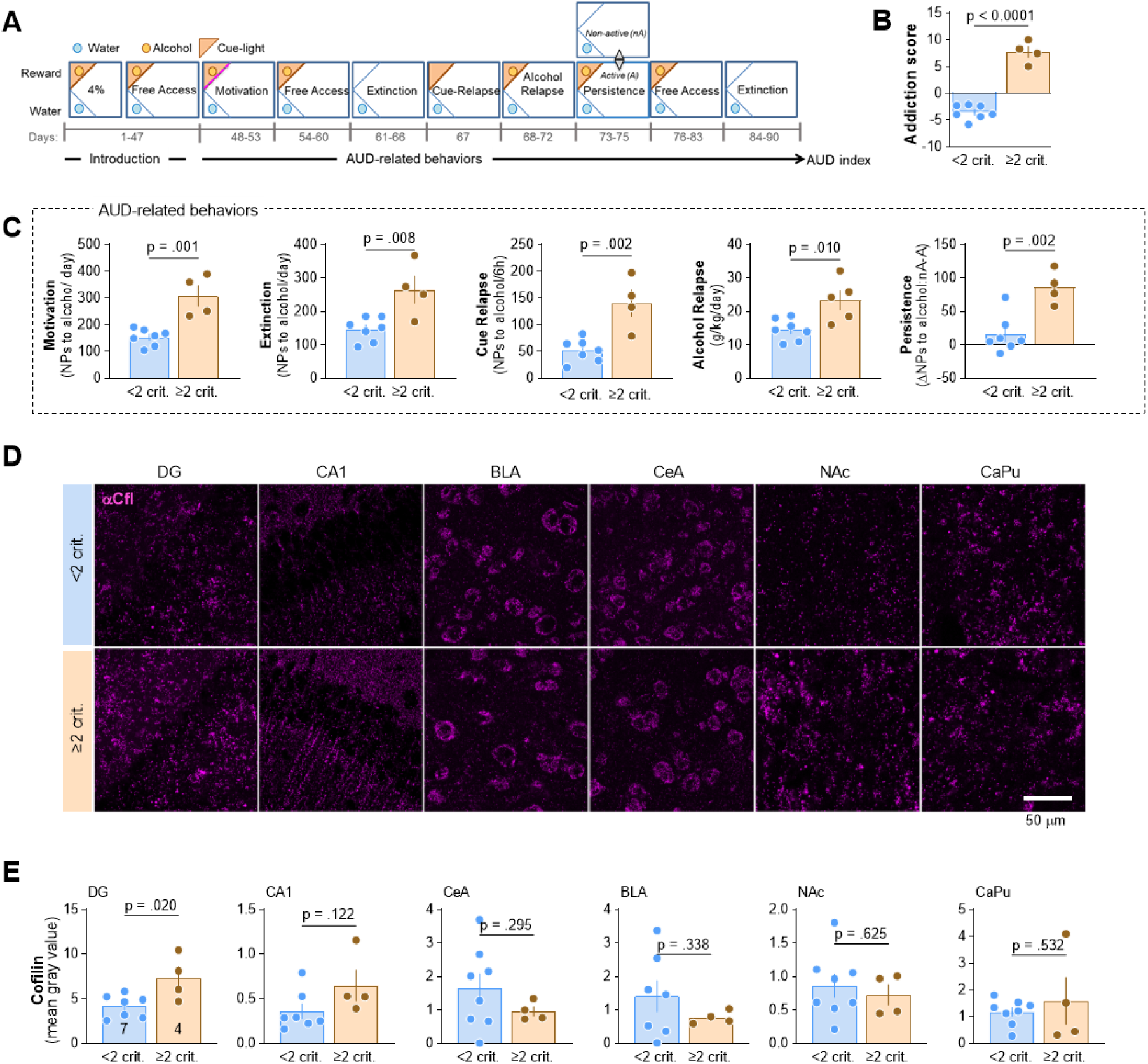
Cfl protein levels are upregulated during alcohol withdrawal in DG in ≥ 2 crit vs 2 crit mice. **(A)** IntelliCage setup and experimental timeline. Mice were trained to drink alcohol (4% and 8%) and AUD-related behaviors were tested: motivation to drink alcohol (Motivation), alcohol seeking during withdrawal (Withdrawal) and cue relapse (Cue relapse), alcohol drinking during alcohol relapse (Alcohol relapse), and alcohol seeking during a persistence test (Persistence). During periods of free access to alcohol (FA) mice had unlimited access to alcohol (8%). ≥ 2 crit and < 2 crit mice were identified based on AUD index (AI). Mice were sacrificed after alcohol withdrawal (withdrawal, day 90). **(B)** AUD index was calculated as a sum of normalized scores in AUD tests. **(C)** Assessment of AUD-related behaviors in ≥ 2 crit and < 2 crit mice: motivation towards alcohol (t(9) = 4.76), extinction of alcohol seeking during withdrawal (t(9) = 3.42) and cue-induced relapse (t(9) = 4.27), alcohol consumption during alcohol relapse (t(9) = 3.15), and persistence in alcohol seeking (t(9) = 4.20). **(D-E)** Analysis of cofilin immunostaining in DG, CA1, basolateral amygdala (BLA), central nucleus of the amygdala (CeA), nucleus accumbens (NAc) and caudate putamen (CaPu). **(D)** Representative microphotographs and **(E)** summary of data (DG: t(9) = 2.82; CA1: t(9) = 1.71; BLA: t(9) = 1.01; CeA: t(9) = 1.11; NAc: t(9) = 0.505; CaPu: t(9) = 0.647).

**Supplementary figure 4.**
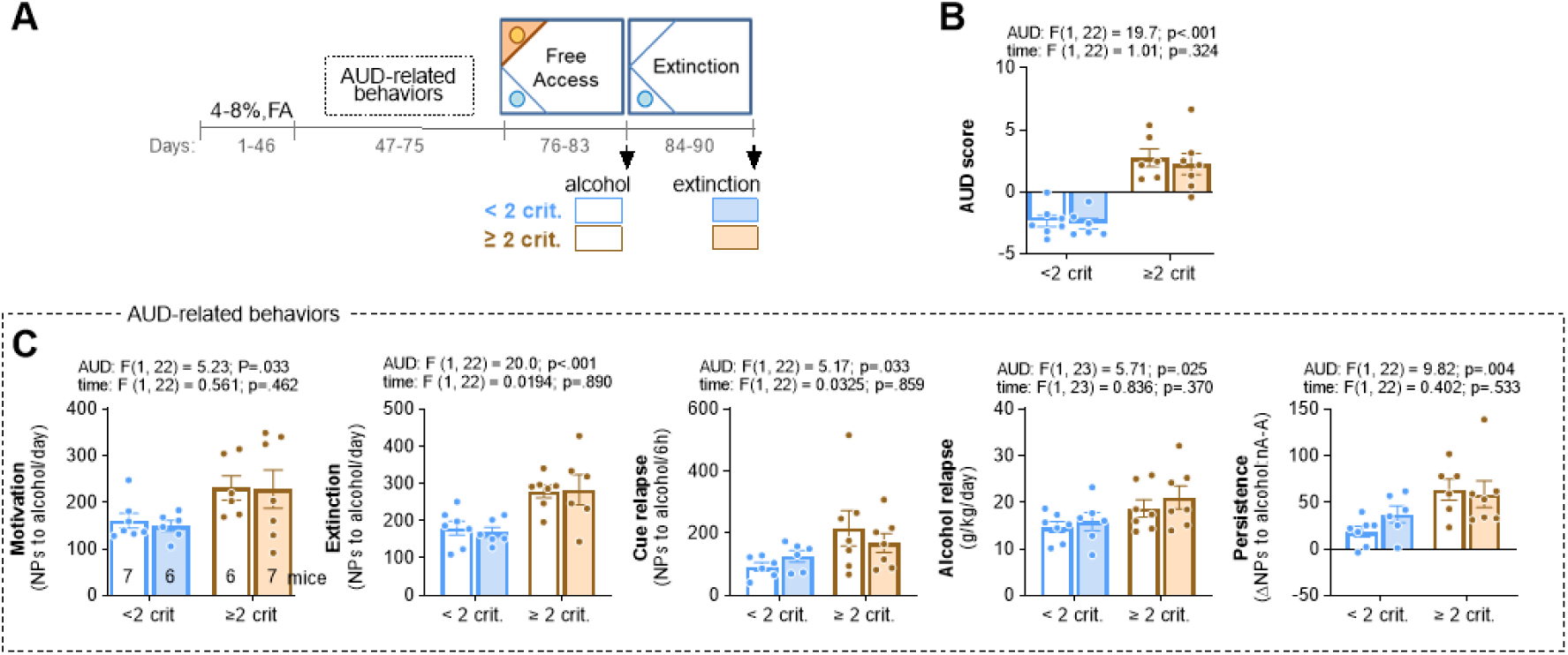
Behavioral characteristics of the < 2 crit and ≥ 2 crit mice (immunofluorescent study). **(A)** IntelliCage setup and experimental timeline. Mice (n = 28) were trained to drink alcohol (4% and 8%) and AUD-related behaviors were tested: motivation to drink alcohol (Motivation), alcohol seeking during withdrawal (Withdrawal) and cue relapse (Cue relapse), alcohol drinking during alcohol relapse (Alcohol relapse), and alcohol seeking during a persistence test (Persistence). During periods of free access to alcohol (FA) mice had unlimited access to alcohol (8%). ≥ 2 crit and < 2 crit mice were identified based on AUD index (AI). Mice were sacrificed after alcohol free access period (alcohol, day 83) or after alcohol withdrawal (withdrawal, day 90). **(B)** AUD index was calculated as a sum of normalized scores in AUD tests. **(C)** Assessment of AUD-related behaviors in ≥ 2 crit and < 2 crit mice: motivation towards alcohol, extinction of alcohol seeking during withdrawal and cue-induced relapse, alcohol consumption during alcohol relapse, and persistence in alcohol seeking (two-way ANOVA tests were used, effects of phenotype (AUD) and time are shown on graphs).

**Supplementary figure 5.**
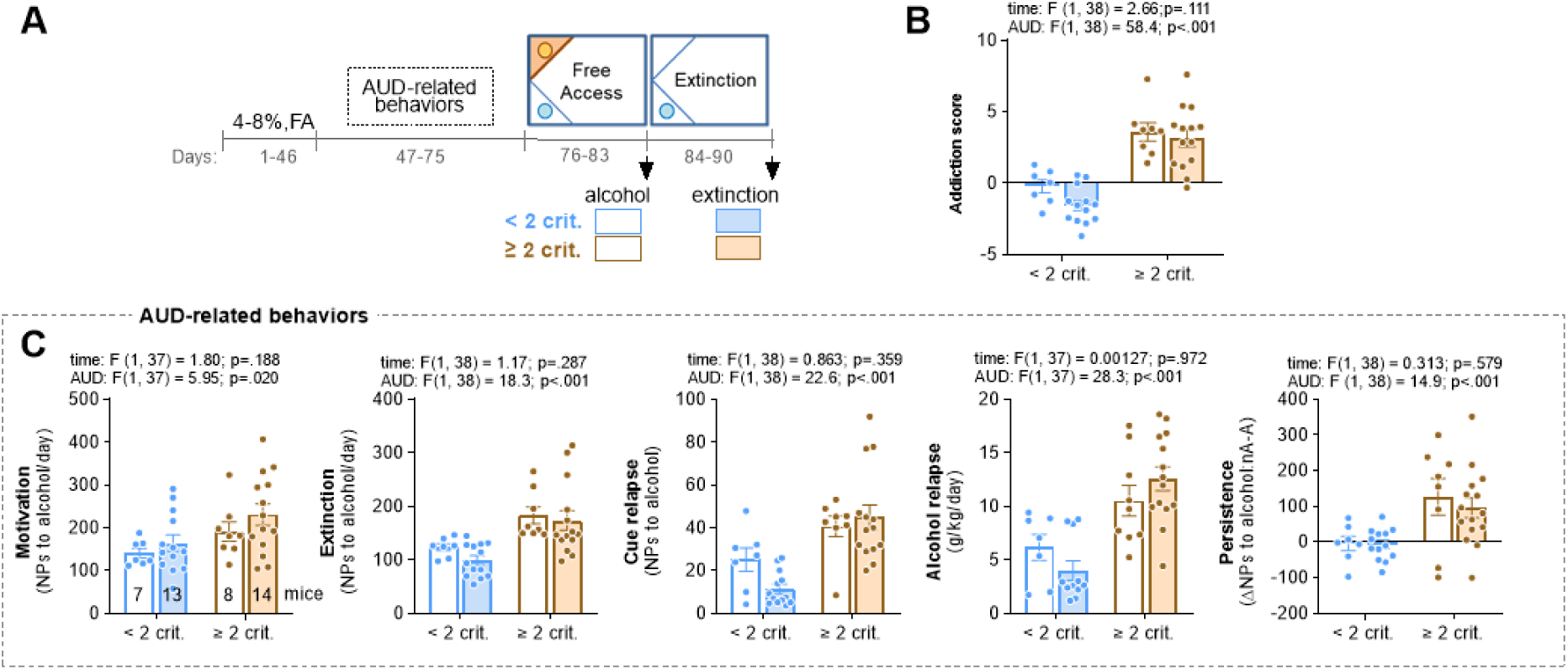
Behavioral characteristics of the < 2 crit and ≥ 2 crit mice (ephys study). **(A)** IntelliCage setup and experimental timeline. Mice were trained to drink alcohol (4% and 8%) and AUD-related behaviors were tested: motivation to drink alcohol (Motivation), alcohol seeking during withdrawal (Withdrawal) and cue relapse (Cue relapse), alcohol drinking during alcohol relapse (Alcohol relapse), and alcohol seeking during a persistence test (Persistence). During periods of free access to alcohol (FA) mice had unlimited access to alcohol (8%). ≥ 2 crit and < 2 crit mice were identified based on AUD index (AI). Mice were sacrificed after alcohol free access period (alcohol, day 83) or after alcohol withdrawal (withdrawal, day 90). **(B)** AUD index was calculated as a sum of normalized scores in AUD tests (two-way ANOVA test was used, effects of AUD index and time are shown). **(C)** Assessment of AUD-related behaviors in ≥ 2 crit and < 2 crit mice: motivation towards alcohol, extinction of alcohol seeking during withdrawal and cue-induced relapse, alcohol consumption during alcohol relapse, and persistence in alcohol seeking (two-way ANOVA tests were used, effects of phenotype (AUD index) and time are shown on the graphs).

**Supplementary figure 6.**
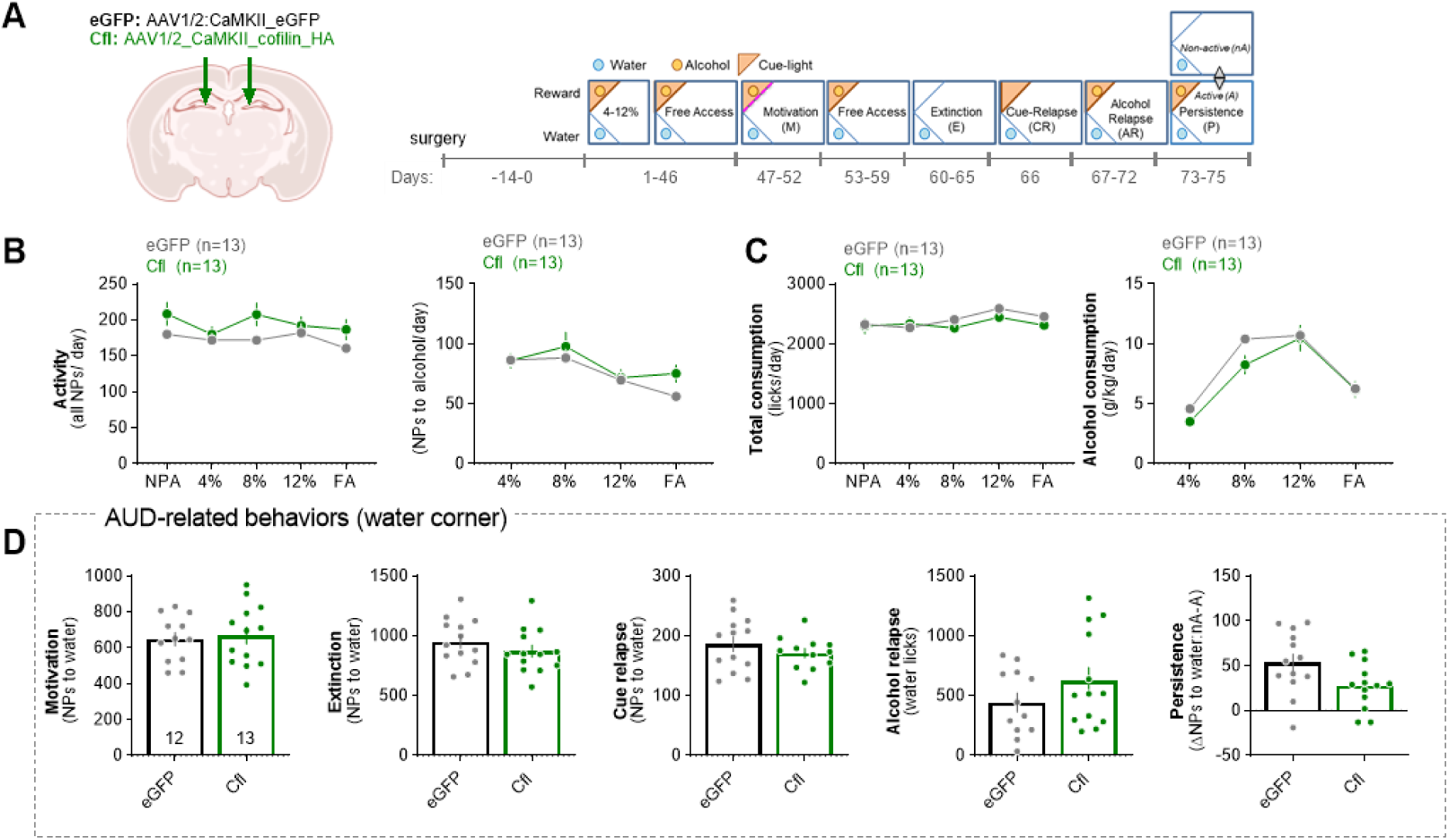
Overexpression of Cfl in PoDG does not affect general activity and animal behavior in the water corner. **(A)** Experimental timelines and IntelliCage setups. Mice received dDG-targeted bilateral stereotactic injections of AAV2.1 encoding cofilin-HA (n = 13), or eGFP (n = 13). Mice were trained to drink alcohol (4-12% and Free access) and AUD-related behaviors were tested: motivation to drink alcohol (M), alcohol seeking during withdrawal (W) and cue relapse (CR), alcohol drinking during alcohol relapse (AR), and alcohol seeking during a persistence test (P). **(B-C)** General activity and consumption. Summary of data showing: **(B)** all nosepokes (repeated measure ANOVA, effect of time: F(2.66, 63.) = 1.98, p = 0.133; effect of virus: F(1, 24) = 2.47, p = 0.129), nosepokes to alcohol (effect of time: F(2.32, 51.1) = 15.7, p < 0.001; effect of virus: F(1, 22) = 0.745, p = 0.397), **(H)** total liquid consumption (effect of time: F(2.55, 28.0) = 3.61, p = 0.031; effect of virus: F(1, 11) = 0.465, p = 0.510) and alcohol consumption (effect of time: F(2.34, 58.4) = 65.5, p < 0.001; effect of virus: F(1, 25) = 1.28, p = 0.269). **(F)** Summary of data showing mice activity in the water corner during AUD-related tests: motivation (t(23) = 0.319, p = 0.753), extinction (t(23) = 1.04, p = 0.309), cue relapse (t(23) = 1.00, p = 0.327), water consumption during alcohol relapse (t(23) = 1.33, p = 0.198) and persistence test (t(23) = 2.15, p = 0.042). Mean ± SEM are shown. Each dot on the D graphs represents one animal.

## SUPPLEMENTARY TABLES

**Supplementary Table 1.** Characteristics of each factor for the factor analysis of ≥ 2 crit. mice. Two factors were extracted (eigenvalue > 1). % tot var, percentage of total variation explained by each factor; Cumul %, cumulative percentage.

| PC summary | Eigenvalue | % total var | Cumul % |
| --- | --- | --- | --- |
| **PC1** | 2,262 | 45,25% | 45,25% |
| **PC2** | 1,160 | 23,20% | 68,44% |
| **PC3** | 0,873 | 17,46% | 85,91% |
| **PC4** | 0,395 | 7,90% | 93,81% |
| **PC5** | 0,310 | 6,19% | 100,00% |

**Supplementary Table 2.** Score of factor loadings of each variable in the factor analysis for ≥ 2 crit. mice. Variables correspond to the following parameters: motivation to drink alcohol (M: a number of nose-pokes in the reward corner performed in a progressive-ratio schedule of reinforcement test when mice had to make an increasing number of nosepokes (FR2, 4, 8, 12, 16, 20, 24, 28…) in order to get access to alcohol for 5 seconds; extinction of alcohol seeking during protracted abstinence (E: a number of nosepokes in the reward corner when the reward corner was inactive and nosepokes had no programmed consequences); reactivity to alcohol-predicting cues (CR: as nosepokes in the reward corner during presentation of the cue light when alcohol was not available) ^43^; lack of control over alcohol consumption when the alcohol corner was activated (AR: g/kg/day); persistence in alcohol seeking, even during signaled alcohol non-availability (P: a change of nosepokes number to the alcohol corner during the non-active vs. active phases of the test).

| Var | PC1 | PC2 | PC3 |
| --- | --- | --- | --- |
| **E** | 0,809 | 0,264 | 0,332 |
| **P** | 0,806 | 0,039 | 0,445 |
| **CR** | 0,638 | -0,353 | -0,584 |
| **AR** | 0,632 | 0,490 | -0,463 |
| **M** | 0,388 | -0,851 | 0,099 |

**Supplementary Table 3.**
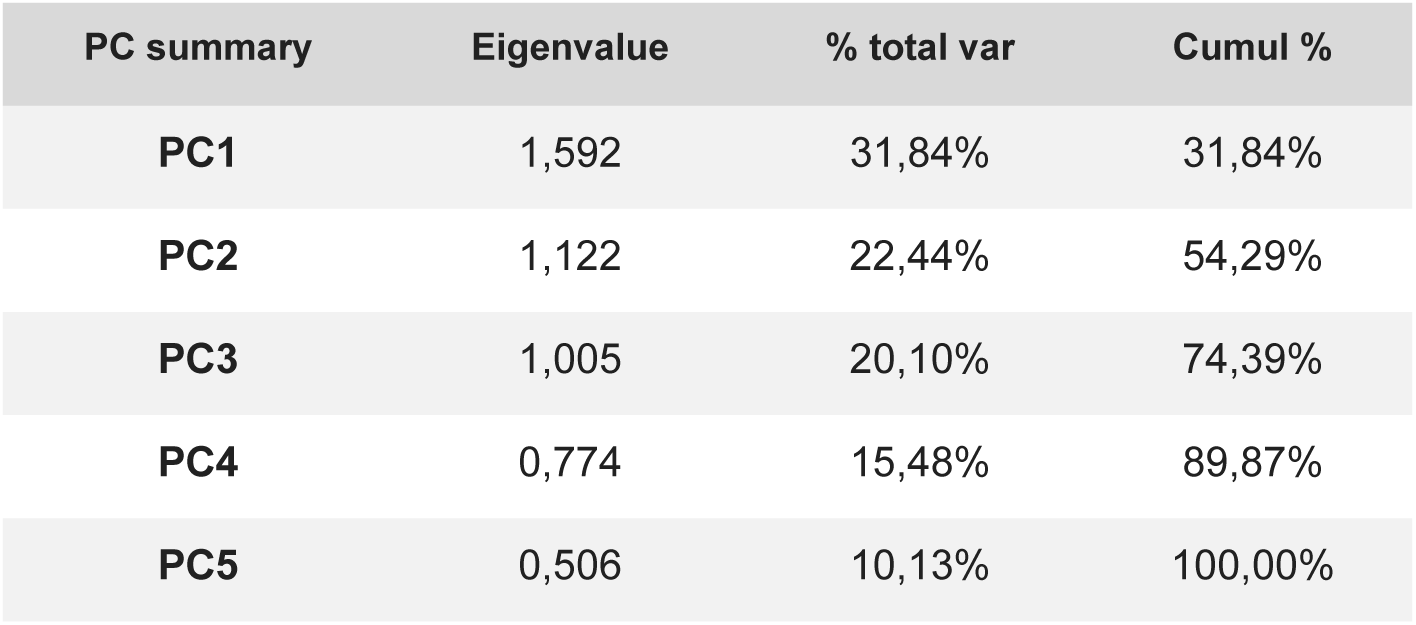
Characteristics of each factor for the factor analysis of < 2 crit. mice. Two factors were extracted (eigenvalue > 1). % tot var, percentage of total variation explained by each factor; Cumul %, cumulative percentage.

**Supplementary Table 4.** Score of factor loadings of each variable in the factor analysis for < 2 crit. mice. Variables correspond to the following parameters: motivation to drink alcohol (M: a number of nose-pokes in the reward corner performed in a progressive-ratio schedule of reinforcement test when mice had to make an increasing number of nosepokes (FR2, 4, 8, 12, 16, 20, 24, 28…) in order to get access to alcohol for 5 seconds; extinction of alcohol seeking during protracted abstinence (E: a number of nosepokes in the reward corner when the reward corner was inactive and nosepokes had no programmed consequences); reactivity to alcohol-predicting cues (CR: as nosepokes in the reward corner during presentation of the cue light when alcohol was not available) ^43^; lack of control over alcohol consumption when the alcohol corner was activated (AR: g/kg/day); persistence in alcohol seeking, even during signaled alcohol non-availability (P: a change of nosepokes number to the alcohol corner during the non-active vs. active phases of the test).

| Var | PC1 | PC2 | PC3 | PC4 |
| --- | --- | --- | --- | --- |
| **M** | 0,828 | -0,132 | 0,110 | -0,170 |
| **P** | 0,243 | 0,247 | 0,914 | 0,125 |
| **CR** | 0,139 | -0,892 | 0,150 | -0,306 |
| **E** | -0,633 | -0,463 | 0,267 | 0,443 |
| **AR** | -0,654 | 0,183 | 0,253 | -0,663 |

**Supplementary Table 5.**
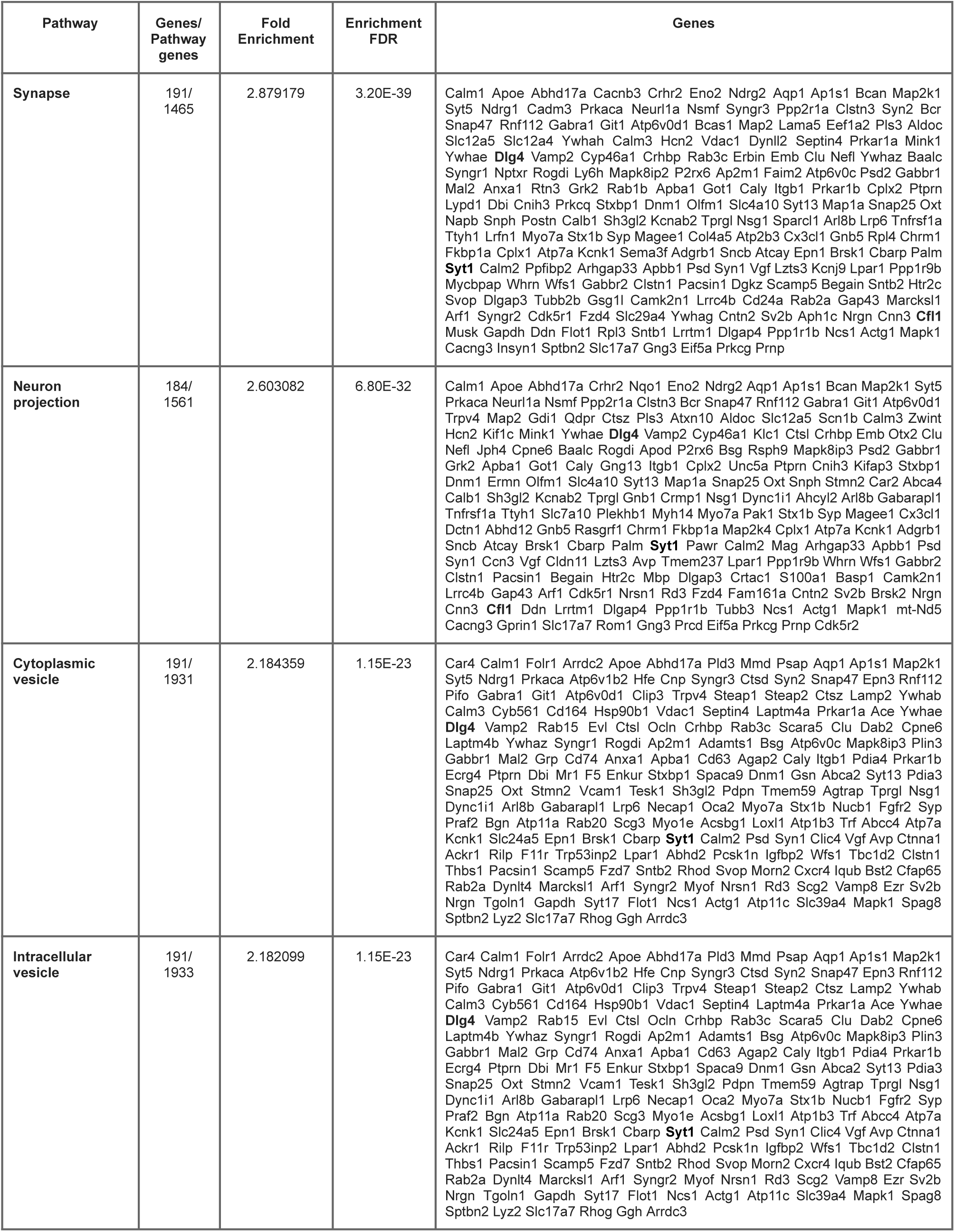
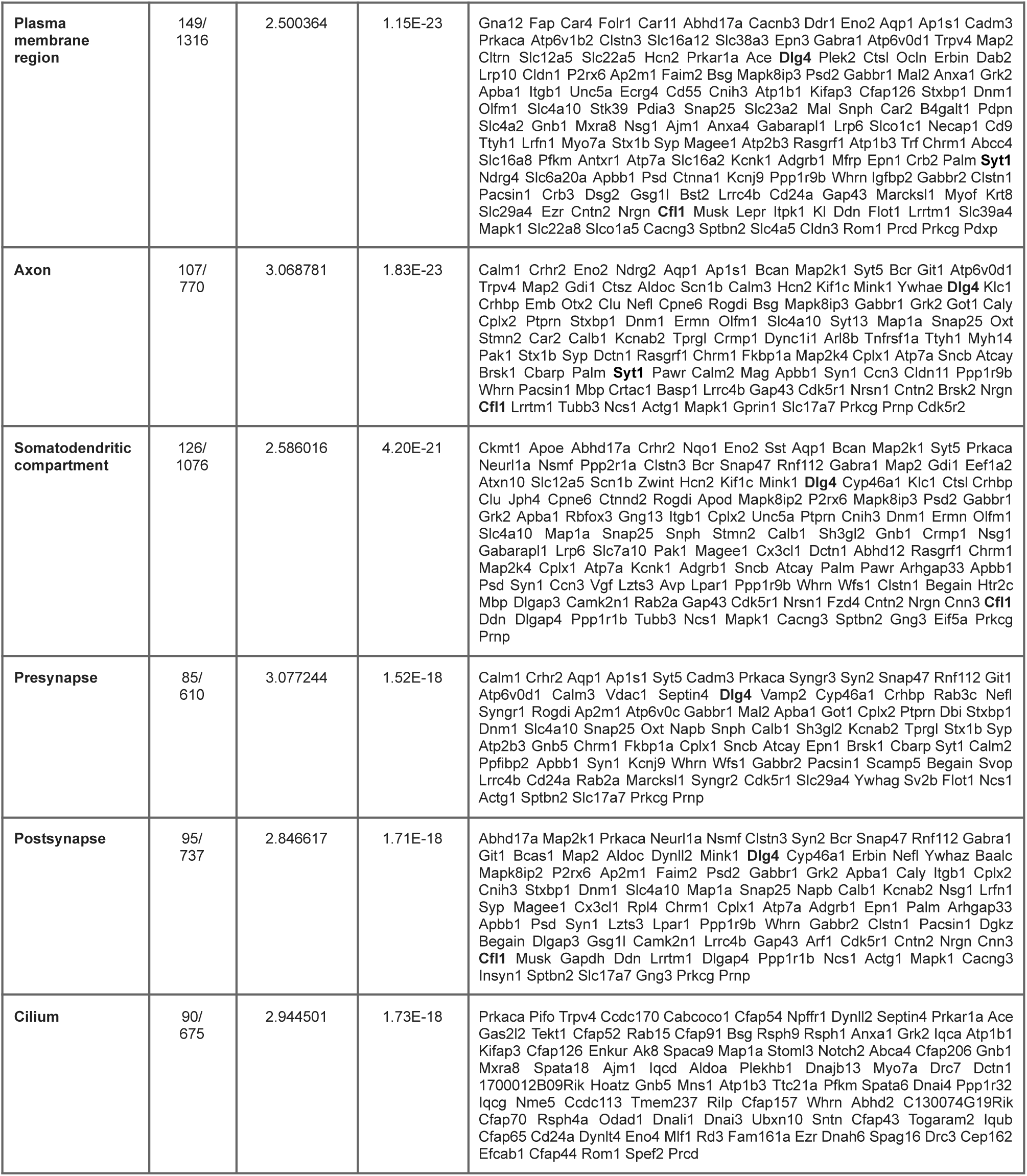
HIPPOCAMPUS, < 2 crit vs ≥ 2 crit mice, GO: Cellular component.

**Supplementary Table 6.**
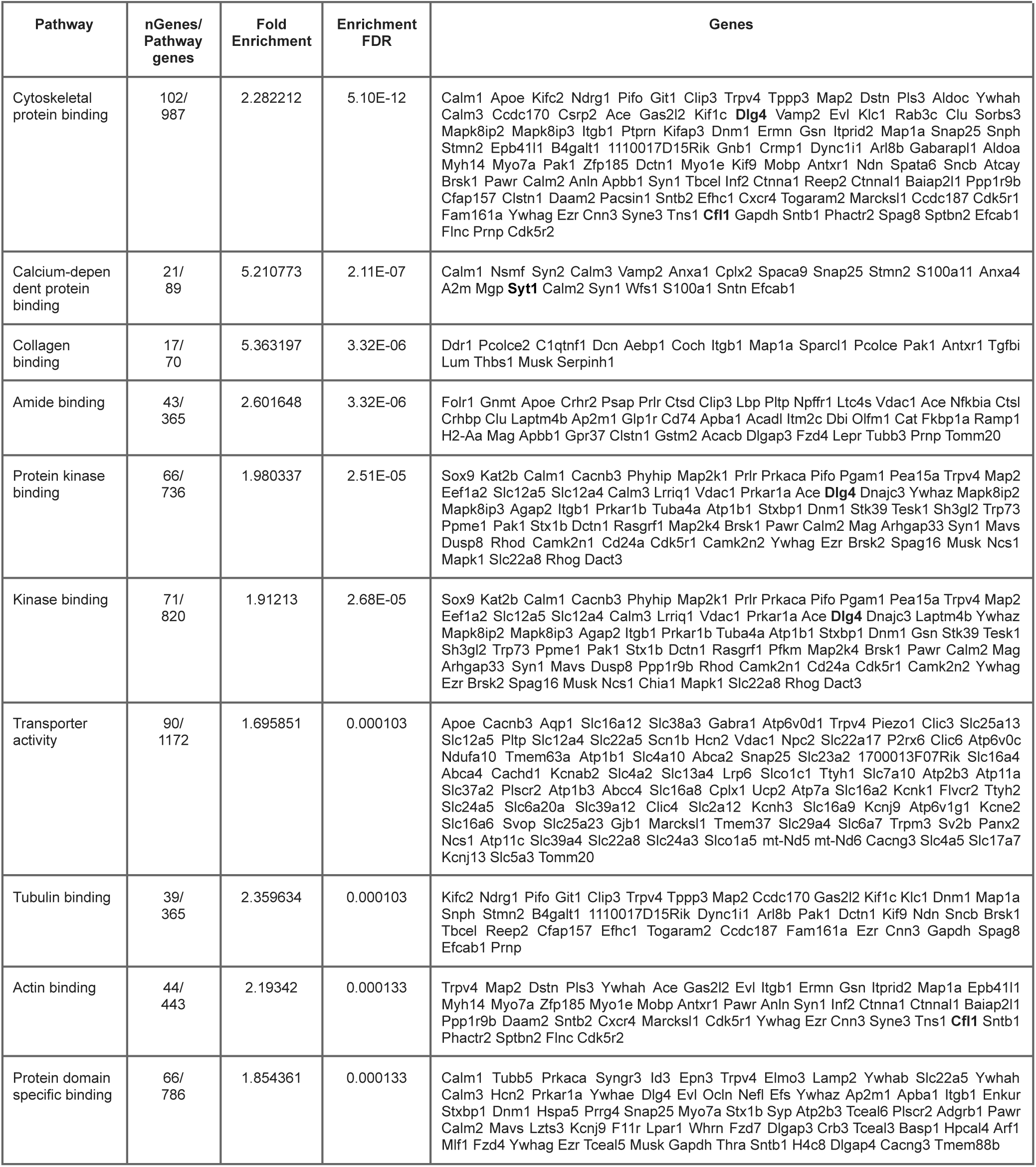
HIPPOCAMPUS, < 2 crit vs ≥ 2 crit mice, GO: Molecular function.

**Supplementary Table 7.**
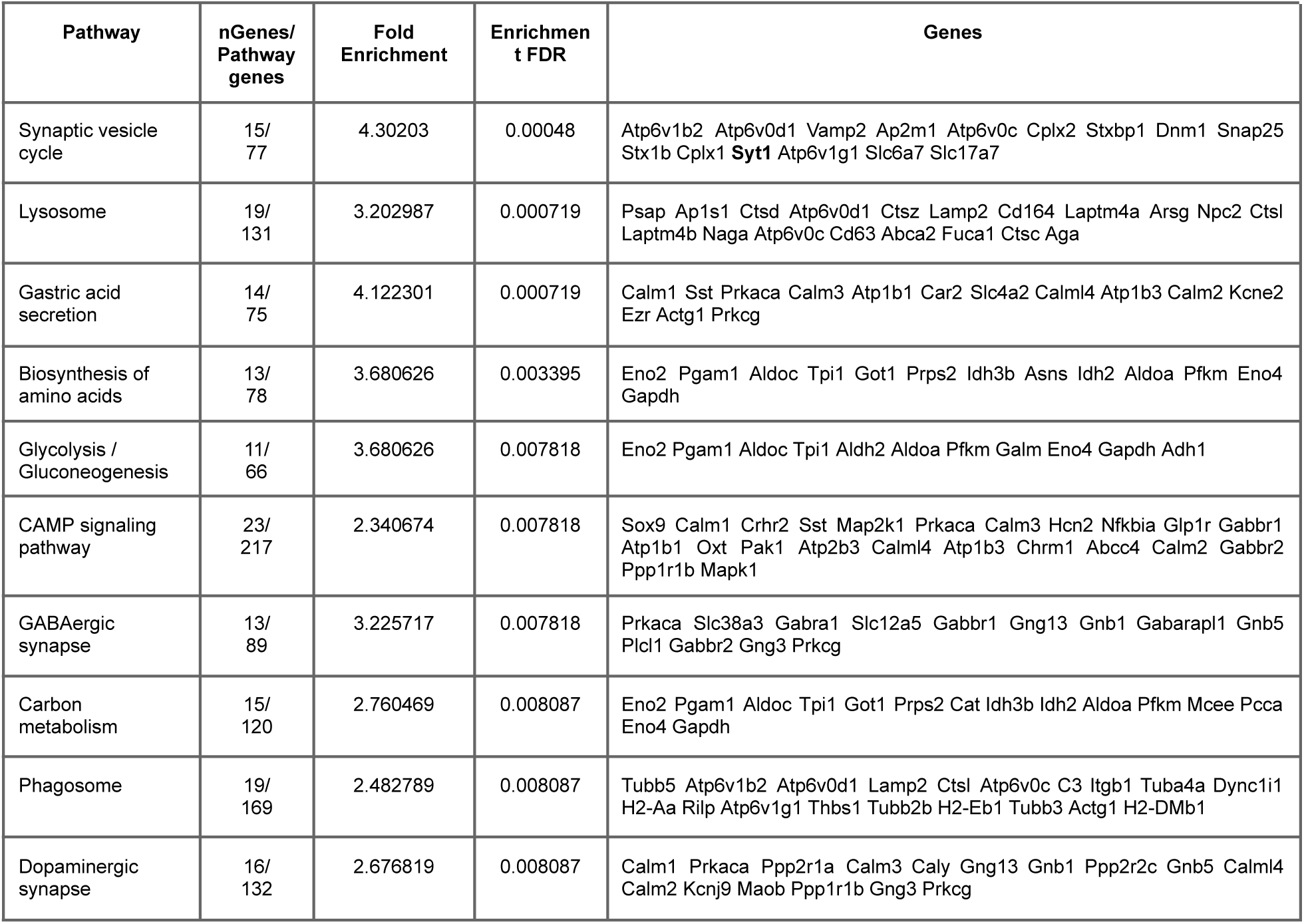
HIPPOCAMPUS, < 2 crit vs ≥ 2 crit mice, KEGG pathways.

**Supplementary Table 8.**
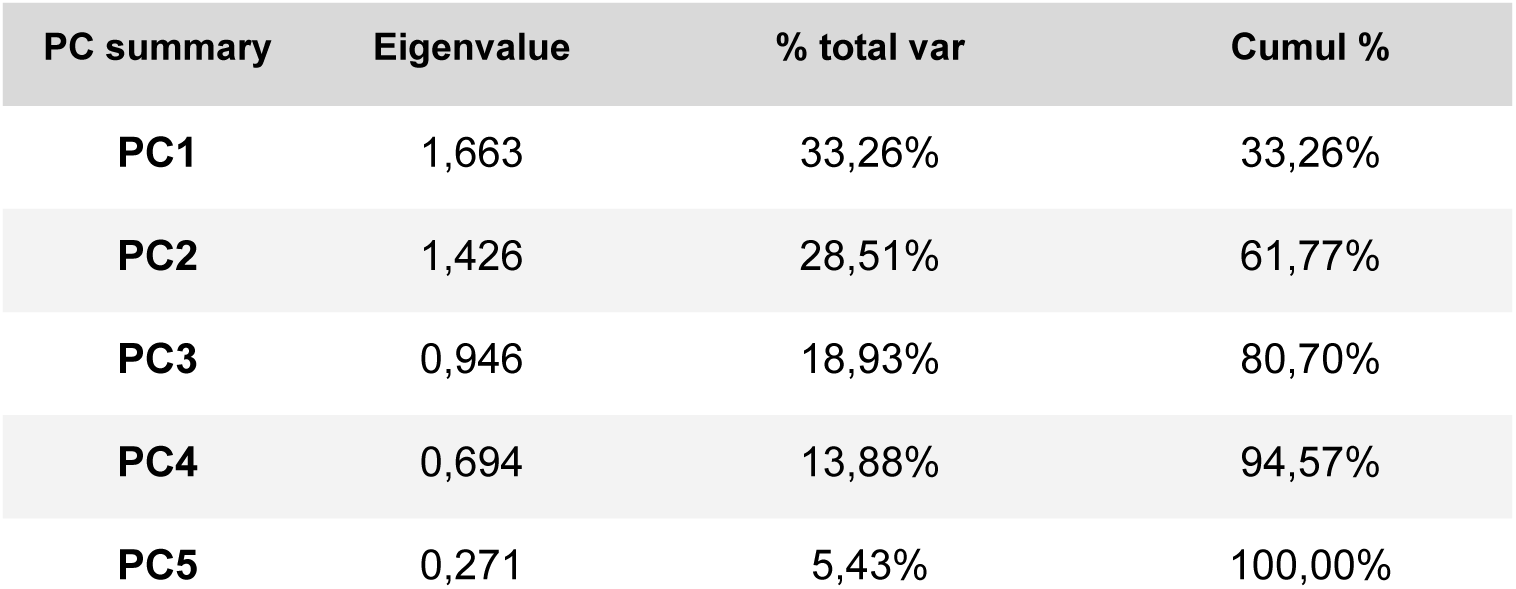
Characteristics of each factor for the factor analysis of eGFP mice. Two factors were extracted (eigenvalue > 1). % tot var, percentage of total variation explained by each factor; Cumul %, cumulative percentage.

**Supplementary Table 9.** Score of factor loadings of each variable in the factor analysis for eGFP mice. Variables correspond to the following parameters: motivation to drink alcohol (M: a number of nose-pokes in the reward corner performed in a progressive-ratio schedule of reinforcement test when mice had to make an increasing number of nosepokes (FR2, 4, 8, 12, 16, 20, 24, 28…) in order to get access to alcohol for 5 seconds; extinction of alcohol seeking during protracted abstinence (E: a number of nosepokes in the reward corner when the reward corner was inactive and nosepokes had no programmed consequences); reactivity to alcohol-predicting cues (CR: as nosepokes in the reward corner during presentation of the cue light when alcohol was not available) ^43^; lack of control over alcohol consumption when the alcohol corner was activated (AR: g/kg/day); persistence in alcohol seeking, even during signaled alcohol non-availability (P: a change of nosepokes number to the alcohol corner during the non-active vs. active phases of the test).

| Var | PC1 | PC2 |
| --- | --- | --- |
| **AR** | 0,741 | -0,157 |
| **M** | 0,247 | 0,850 |
| **E** | -0,448 | -0,560 |
| **CR** | -0,520 | 0,603 |
| **P** | -0,763 | 0,040 |

**Supplementary Table 10.**
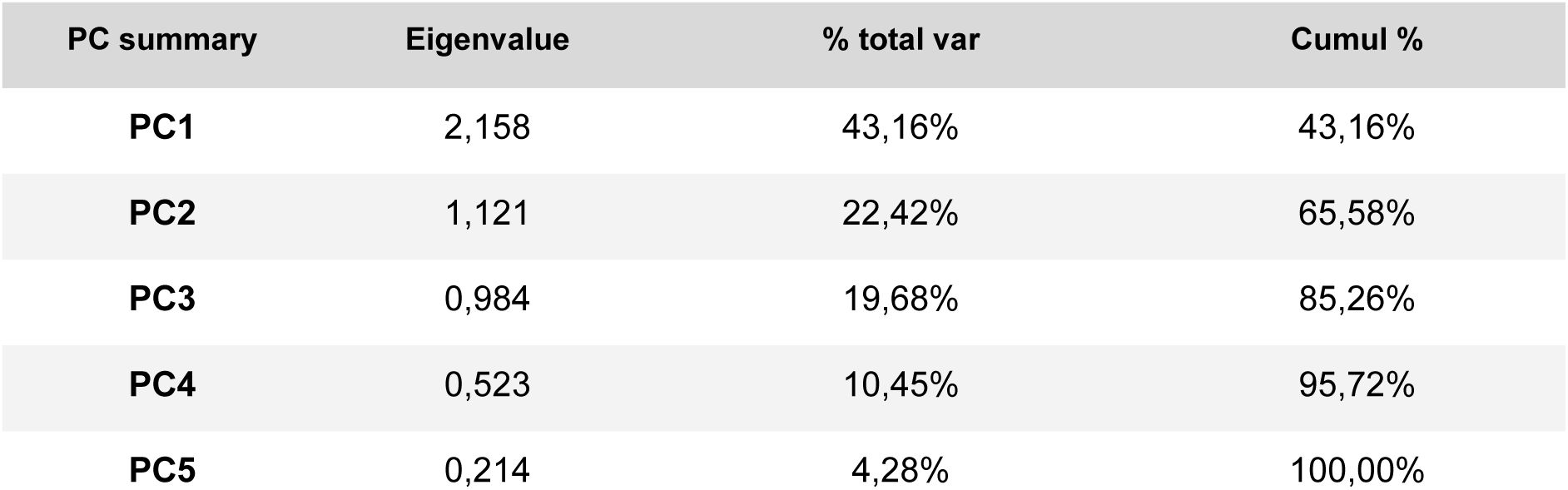
Characteristics of each factor for the factor analysis of Cfl mice. Two factors were extracted (eigenvalue > 1). % tot var, percentage of total variation explained by each factor; Cumul %, cumulative percentage.

| PC summary | Eigenvalue | % total var | Cumul % |
| --- | --- | --- | --- |
| **PC1** | 2,158 | 43,16% | 43,16% |
| **PC2** | 1,121 | 22,42% | 65,58% |
| **PC3** | 0,984 | 19,68% | 85,26% |
| **PC4** | 0,523 | 10,45% | 95,72% |
| **PC5** | 0,214 | 4,28% | 100,00% |

**Supplementary Table 11.** Score of factor loadings of each variable in the factor analysis for Cfl mice. Variables correspond to the following parameters: motivation to drink alcohol (M: a number of nose-pokes in the reward corner performed in a progressive-ratio schedule of reinforcement test when mice had to make an increasing number of nosepokes (FR2, 4, 8, 12, 16, 20, 24, 28…) in order to get access to alcohol for 5 seconds; extinction of alcohol seeking during protracted abstinence (E: a number of nosepokes in the reward corner when the reward corner was inactive and nosepokes had no programmed consequences); reactivity to alcohol-predicting cues (CR: as nosepokes in the reward corner during presentation of the cue light when alcohol was not available) ^43^; lack of control over alcohol consumption when the alcohol corner was activated (AR: g/kg/day); persistence in alcohol seeking, even during signaled alcohol non-availability (P: a change of nosepokes number to the alcohol corner during the non-active vs. active phases of the test).

| Var | PC1 | PC2 | PC3 |
| --- | --- | --- | --- |
| **AR** | 0,882 | -0,114 | 0,305 |
| **E** | 0,728 | -0,461 | -0,102 |
| **M** | 0,674 | 0,475 | 0,465 |
| **P** | 0,568 | -0,175 | -0,685 |
| **CR** | 0,27 | 0,799 | -0,442 |

